# Novel organelle anion channels formed by chromogranin B drive normal granule maturation in endocrine cells

**DOI:** 10.1101/302828

**Authors:** Gaya P. Yadav, Hui Zheng, Qing Yang, Lauren G. Douma, Mani Annamalai, Linda B. Bloom, Clayton Mathews, Qiu-Xing Jiang

## Abstract

All endocrine cells need an anion conductance for maturation of secretory granules. Identity of this family of anion channels has been elusive for forty years. We now show that a family of granule protein, CHGB, serves the long-sought conductance. CHGB interacts with membranes through two amphipathic helices, and forms a chloride channel with large conductance and high anion selectivity. Fast kinetics and high cooperativity suggest that CHGB tetramerizes to form a functional channel. Nonfunctional mutants separate CHGB’s function in granule maturation from that in granule biogenesis. In neuroendocrine cells, CHGB channel and a H^+^-ATPase drives normal insulin maturation inside or dopamine loading into secretory granules. CHGB’s tight membrane-association after exocytotic release of secretory granules separates its intracellular function from extracellular functions of its proteolytic peptides. CHGB-null mice show consistent impairment of granule acidification in pancreatic beta-cells. These findings together support that the phylogenetically conserved CHGB proteins constitute a new family of organelle chloride channels in the regulated secretory pathway among various endocrine cells.

## INTRODUCTION

Cells rely on secretory pathways to send out specific bioactive molecules to their surroundings (Grimes and Kelly, 1992). Both constitutive and regulated secretory pathways have been studied extensively since early EM observations of intracellular vesicular trafficking. Many molecular players in these pathways are known, and conserved mechanisms for vesicle targeting and membrane fusion are established (Farquhar and Wellings, 1957; Jamieson and Palade, 1964; Rizo and Xu, 2015; Rothman, 1994; Salama and Schekman, 1995). Three key steps of regulated secretion are granule biogenesis at the *trans* Golgi network (TGN), granule maturation through fusion and membrane shedding, and exocytotic release of mature granules (Hosaka and Watanabe, 2010). It has been long known that maturation of nascent immature secretory granules (ISGs) into dense-core granules (DCGs) needs a vesicular H^+^-ATPase and a Cl^−^ conductance (Johnson et al., 1982; Johnson and Scarpa, 1976). Till now, identity of the Cl^−^ conductance remains uncertain. ClC-3 Cl^−^/H^+^ co-transporter was once proposed as a candidate, but its granular localization was ultimately negated (Deriy et al., 2009; Jentsch et al., 2010; Li et al., 2009; Maritzen et al., 2008). Further, ClC-3’s fS conductance and unmeasurable outward flux make it unfit to serve the unknown Cl^−^ conductance for granule acidification (Fahlke, 2001; Guzman et al., 2013). Such a missing link has limited our understanding of granule acidification and cargo maturation.

Granin family proteins are by default secretory proteins that chaperone other secretory molecules through regulated secretion (Bartolomucci et al., 2011; Hosaka and Watanabe, 2010). They have intracellular functions and extracellular roles. Chromogranins (CHGs) are believed to function in all three steps of the pathway. It was proposed that they interact with cargos, serve as low-affinity, high-capacity Ca^2+^ reserve. Their extracellular functions are executed by CHG-derived peptides that are associated with various human diseases (Bartolomucci et al., 2011). However, the molecular mechanisms for all CHGs’ intracellular functions remain to be elucidated.

Members of the granin superfamily (Bartolomucci et al., 2011) share low amino acid sequence similarity (Montero-Hadjadje et al., 2008) and are clustered into phylogenetic subfamilies. Granin proteins of distinct subfamilies usually coexist in secretory granules and were hypothesized to work with different partners (Bartolomucci et al., 2011). Native CHGB forms high-order aggregates at low pH and with mM Ca^2+^ (Yoo, 1995a, b). Partially purified native CHGB associates quite strongly with lipid vesicles (Yoo, 1995b). A “tightly membrane-associated form” of CHGB can be observed on the surface of PC-12 cells after stimulated granule release (Pimplikar and Huttner, 1992). CHGB is genetically associated with neurodegeneration and psychiatric disorders (Davenport et al., 2010; Gros-Louis et al., 2009). It was proposed to participate in forming TGN proteineous aggregates and driving granule biogenesis (Takeuchi and Hosaka, 2008; Tooze, 1998). Studies of CHGB–*null* mice by two groups reported conflicting results regarding CHGB’s role in granule biogenesis (Diaz-Vera et al., 2010; Obermuller et al., 2010; Zhang et al., 2014; Zhang et al., 2009). A mechanistic view on CHGB’s intracellular functions is still unavailable.

In this work we provide the first elucidation of an important intracellular function of CHGB in the regulated secretory pathway. Starting from the contradiction in literature between the CHGB being tightly membrane-bound (Pimplikar and Huttner, 1992; Yoo, 1995b) and its isolation partially in heat-stable fractions or soluble protein complexes, we investigate the CHGB-membrane interaction in vitro, in culture cells, and in knockout mice to reveal a novel chloride channel function of the CHGB that is needed for normal granule maturation in endocrine cells and thus in a perfect location to serve the long-sought anion conductance first proposed four decades ago.

## RESULTS

### CHGB inserts itself into membranes and induces nanotubules from bilayers

To study CHGB function, recombinant murine CHGB was purified from *sf9* cells. During biochemical preparation, Triton X-100-like detergents were needed to keep CHGB soluble. In size-exclusion chromatography (SEC), purified CHGB was eluted as a single, symmetric peak (Fig. 1A) with a size equivalent to an ∼300 kDa globular protein. Due to posttranslational modifications, a high content of charged residues, or possibly detergent-binding, the recombinant CHGB ran at ∼86 kDa in a reducing SDS-PAGE gel (Fig. 1B, 1C), which is larger than the 78 kDa calculated from its sequence, behaving similarly to mature human CHGB (Pimplikar and Huttner, 1992). Due to the detergent micelle (∼100 kDa), the CHGB was likely a dimer, instead of a trimer. The detergent-solubilized CHGB treated with a bifunctional cross-linker, 4-(N-Maleimidomethyl) cyclohexane-1-carboxylic acid 3-sulfo-N-hydroxysuccinimide ester (sulfo-SMCC), showed cross-linked dimers, trimers, tetramers and high-order oligomers (U_2_, U_3_, U_4_, U_n_ in Fig. 1C), indicating a dynamic equilibrium between dimers and oligomers (U_n_, n >= 4) and dominance of the dimers in detergents. Consistently, small amounts of oligomers were observed in an earlier SEC step during purification (Fig. S1A).

**Figure 1.**
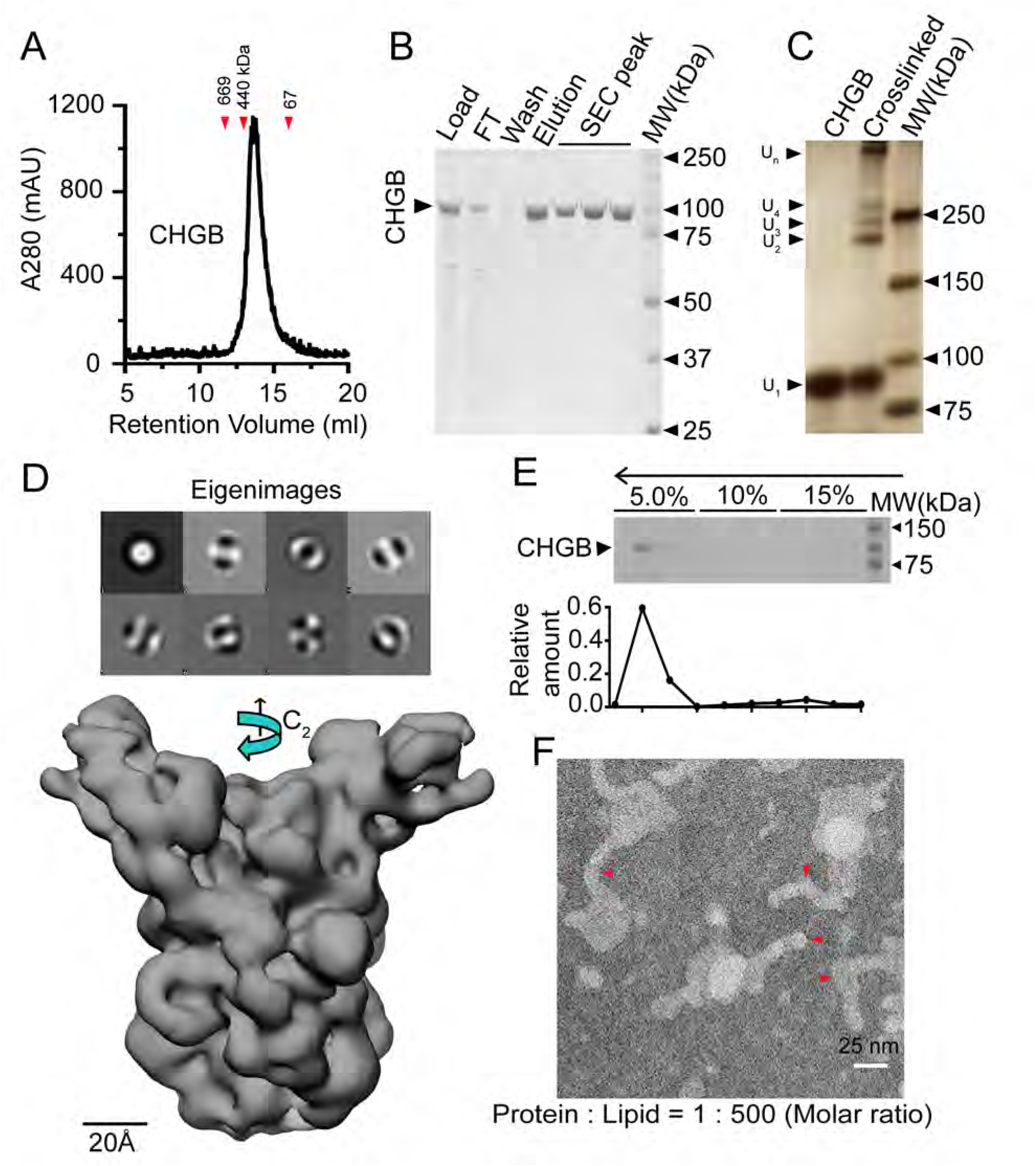
Strong interaction of CHGB and lipid membrane. (**A**) Size-exclusion chromatography (SEC) profile for purified CHGB. Red triangles indicate molecular weight markers. (**B**) SDS-PAGE of CHGB from ion exchange and SEC. (**C**). Silver-stained SDS-PAGE gel of dimers (U_2_), trimers (U_3_), tetramers (U_4_) and high-order oligomers (U_n_) in CHGB treated with Sulfo-SMCC (middle). (**D**) Top: first 8 eigenimages from multivariate statistical analysis of a negative-stain EM dataset show C2 symmetry. Low: a cryoEM map for CHGB dimer at ∼10 Å. (**E**) Top: SDS-PAGE of fractions from a vesicle floatation assay of CHGB in egg PC vesicles in a three-step Ficoll 400 gradient. Top arrow marks the vesicle floatation from bottom (15%) to top (5%). Bottom: Quantified CHGB distribution from SDS-PAGE. (**F**) A typical negative-stain EM image showing nanotubules capped with nanospheres from CHGB vesicles in a PLR of 1 : 500. Red triangles mark areas with strong positive curvature.

Calcium binding is a biochemical hallmark of CHGB due to high content of negatively-charged residues. We examined calcium-induced CHGB aggregates by light-scattering and negative-stain EM (Fig. S1B-F). Fitting of light-scattering data with a Hill equation yielded an apparent *K_d_* ∼0.25 mM for Ca^2+^, and a Hill coefficient of ∼2.0 (Fig. S1B), agreeing with CHGB’s low-affinity and high-capacity for Ca^2+^ (Yoo and Lewis, 1996). EM examination found that without Ca^2+^ CHGB proteins were monodisperse (Fig. S1C). The size and amount of the Ca^2+^-induced aggregates increased significantly when [Ca^2+^] rose from 0.2 to 5.0 mM (Fig. S1D-F).

To resolve the ambiguity between CHGB dimers and trimers in detergents, we performed single particle reconstruction. More than 5,400 images of negatively stained CHGB molecules (Fig. S1G) were assembled for multi-variate statistical analysis (MSA). The eigenimages from MSA (top in Fig. 1D) showed strong C2 symmetry. Angular reconstitution and 3D refinement yielded a negative-stain map at ∼30 Å (Fig. S1H). CHGB’s small mass makes it difficult to visualize under cryoEM. We used a ChemiC (chemically functionalized carbon) method (Llaguno et al., 2014) to enhance cryoEM image contrast (Fig. S1I). More than 24,000 cryoEM images were manually selected for analysis. Multi-rounds of 2D and 3D classification led to a homogenous set of ∼6,900 particles and a 3D map at ∼10 Å resolution (Fig. 1D), estimated by the gold-standard Fourier Shell Correlation. The cryoEM map has C2 symmetry and an elongated shape, explaining its larger apparent size in SEC (Fig. 1A). Its distinct features may correspond to the predicted helical and random-coil regions, and confirm CHGB’s dimeric nature in detergents (Fig. 1A, 1C).

The surprising need of detergents for CHGB extraction intrigued us to study whether it directly interacts with membranes using a vesicle flotation assay (Zheng et al., 2011). After reconstitution with lipids(Lee et al., 2013), CHGB floated with vesicles from bottom to top in a three-step Ficoll 400 gradient (Arrow in the top of Fig. 1E). Quantification of the protein bands revealed that > 98% of CHGB protein was in membrane (Fig. 1E, bottom panel). To visualize the effects of CHGB-membrane interaction on the bilayer structure, we examined CHGB vesicles by negative-stain EM (Fig. 1F, S2A-C). When CHGB : lipid molar ratio (PLR) > 1 : 1,000, equivalent to ∼40 CHGB dimers per 100 nm vesicle, 25 nm nanospheres appeared on vesicles (red arrowheads in Fig S2B). More CHGB led to ∼20 nm-thick nanotubules capped with 25 nm-diameter hemi-nanospheres (Fig. 1F, S2C). In some specimens, the nanospheres were severed into individual soluble nanoparticles. These results reveal CHGB-induced remodeling of membranes when local PLR is high, a condition likely being satisfied at the TGN sites for granule biogenesis, which might contribute to the heat-stable or soluble fraction used in many published studies (Benedum et al., 1986; Benedum et al., 1987). The nanotubules and nanospheres indicate strong positive curvature caused by CHGB. To avoid membrane remodeling, we intentionally limited the PLR < 1: 5,000 in most vesicle-based assays.

Lipid membranes drive CHGB oligomerization. When purified CHGB was first reconstituted into vesicles and then extracted with detergents plus ∼0.1 mg/ml lipids, >70% of CHGB protein was eluted by SEC at a position equivalent to an ∼0.8 MDa globular protein (Fig. S2D), suggesting that lipids may stabilize a high-order oligomer. Due to the elongated shape of the cryoEM map (Fig. 1D), biochemically cross-linked tetramers (Fig. 1C) and the detergent micelles, the smallest lipid-stabilized oligomers may be a tetramer (U_4_; Fig. S1A). It may endow the function of CHGB in membrane.

### Amphipathic segments responsible for strong CHGB-membrane interaction

To examine the biophysical nature of CHGB-induced membrane tubulation, we studied whether CHGB causes membrane leak. A fluorescein-release assay was implemented (Mukherjee et al., 2014) to monitor the increase in fluorescence when the fluorescein was released and unquenched. Our data showed that CHGB vesicles did not leak the 10Å fluorophore (Fig 2A). When CHGB vesicles were loaded with Ca^2+^, a Ca^2+^-sensitive dye, Indo-1, detected no leak (Fig. S3A & 2B). Neither DMSO nor valinomycin triggered fluorescein or Ca^2+^ leak while octyl-beta-glucoside (β-OG) released all (Fig. 2A, 2B). As a positive control, purified IP_3_R vesicles showed IP_3_-triggered Ca^2+^ release (Fig. S3B). We then tested if CHGB vesicles leak Cl^−^ by using a system first described by (Stockbridge et al., 2013). CHGB vesicles containing 300 mM KCl were changed into a buffer with 300 mM K-isethionate plus 0.2 mM KCl before being added into a chamber (diagramed in Fig. S3C). Adding valinomycin, a K^+^ carrier, initiated Cl^−^ leak from vesicles (Fig. 2C). Control vesicles without CHGB showed no signal. These findings suggest that a significant fraction of vesicles released internal Cl^−^.

**Figure 2.**
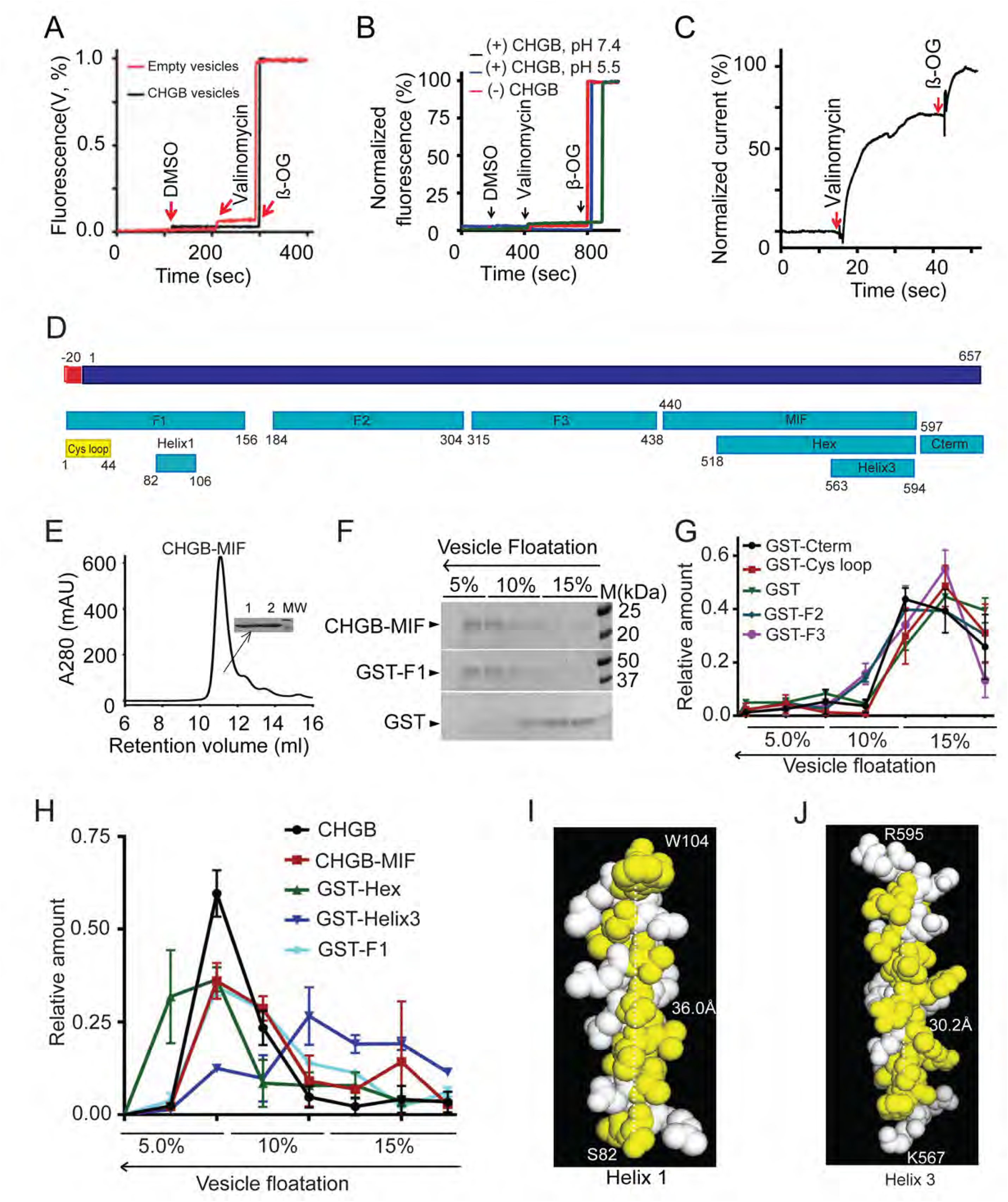
Mapping amphipathic helical segments for CHGB insertion in membrane. (A) CHGB vesicles have no membrane cracks. CHGB vesicles loaded with 1.0 μM carboxy fluorescein were treated with 1.0 μl DMSO, 1.0 μM valinomycin, and finally with 5.0 mM beta-OG to disrupt vesicles. (B) CHGB vesicles do not leak Ca^2+^ at pH 7.4 or 5.5. Vesicles loaded with 1.0 mM CaCl_2_ were treated with 1.0 μl DMSO, 1.0 μM valinomycin, and finally with 5.0 mM beta-OG. (C) Cl^−^ release from vesicles recorded with a Ag/AgCl electrode. 300 mM KCl inside vesicles; 300 mM K-isethionate and 0.2 mM KCl, 10 mM HEPES pH 7.4 outside. 0.25 μM valinomycin started the release. At the end 10 mM β-OG was added to release all Cl^−^. A typical trace out of 3 was shown. Control vesicles did not show valinomycin-triggered response. The recordings were normalized to the β-OG signal. (**D**) A diagram of CHGB and different fragments. (**E**) SEC of CHGB-MIF in a Superdex 200 column. The inset is SDS-PAGE of two peak fractions. (**F**) Vesicle floatation assay for CHGB-MIF, GST-F1 and GST. (**G&H**) Distribution of CHGB and its fragments from vesicle floatation assays as in C & Fig. S3F. Those staying at the bottom in (**G**) and those floating in (**H**). (**I**) A helical model for Helix 1 with its hydrophobic residues in yellow. It was energetically optimized in Coot and presented in PyMol. (**J**) Structural model for Helix 3 (Hex 3).

Chloride leak raised the question of whether CHGB is deeply integrated in membrane. Trypsinization of CHGB vesicles was performed to test this possibility. After 50 min, > 90% CHGB was digested, indicating its preferential insertion from extravesicular side. Trypsin-treatment of Ca^2+^-loaded CHGB vesicles introduced no leak, meaning that trypsinization did not break the membrane (Fig. S3E). Mass-spectrometry and N-terminal sequencing of one membrane interacting fragment (MIF, Fig. S3D) identified CHGB 440-597, named as CHGB-MIF (Fig. 2D). Secondary structure analysis of CHGB-MIF revealed a shorter segment (Hex, CHGB 518-597) containing two α-helices interspaced by a short random-coil loop (Fig. 2D, Helices 2 and 3 in Fig. S4C-D, S4E-F). Recombinant CHGB-MIF alone interacts with membrane (Fig. 2E-F), suggesting that it is a major contributor to CHGB-membrane interaction (Fig. 2C). Preferential insertion from outside is expected to increase outer surface area of membranes and induce positive curvature (Figs 1F & S2B-S2C).

Systematic mapping of MIFs identified two amphipathic helices (helices 1 and 3 in Figs 2D, S4A-G). Based on secondary structure prediction and the CHGB post-processing peptides, we purified multiple short segments as GST-fusion proteins and reconstituted them for vesicle floatation (Figs 2D, 2E-H, Fig. S3F). As a positive control, His-tagged MIF floated well (Figs 2E-F) because it was well protected by membrane (Fig. S3D). Soluble GST alone failed to float (Fig. 2C). Except Helix 3, all fragments (Fig. 2G-H, S3F) fall into two groups: those float and those sink (Fig. 2G vs. 2H). GST-Helix3, the shortest helix, was distributed between the two (Fig. 2H). The N-terminal fragment F1 (Figs 2D, 2H) contains the N-terminal Cys-loop, which is a soluble sorting signal (Cys-loop Fig. 2G), and a long amphipathic helix (Helix 1, Figs 2I, S4A-B, S4E). A structural model in Fig. 2I shows its extensive hydrophobic surface (yellow; ∼36 Å long). Not protected from trypsinization in membranes, Helix 1 probably lies on membrane surface. Helices 2 and 3 (Figs 2J, S4C-D, S4F-G) are well conserved in the CHGB subfamily (Fig. S4D) and have weak and strong hydrophobic moments, respectively (Figs S4F-G). Structural modeling of Helix3 (Fig. 2J) reveals a hydrophobic surface (yellow) that is >30 Å in length, enough to span a typical bilayer. Possible cooperativity within the Hex segment or between Hex and Helix 1 may contribute to the “tightly membrane-associated form” of CHGB (Pimplikar and Huttner, 1992).

### CHGB alone suffices to form an anion-selective channel

Cl^−^ leak from CHGB vesicles suggested either a transporter or a channel (Fig. 2C). We first tested if CHGB is a channel by bilayer recordings (Lee et al., 2013). The membrane-remodeling property made it difficult to record from many CHGB molecules. Instead, when diluted CHGB vesicles of low PLR (<1:10,000) were fused into planar lipid bilayers in the presence of 0.5 mM CaCl_2_, we observed multiple channel events (Fig. 3A). The channels were almost always open in a low voltage range (−50 to +50 mV), and switched off more frequently in a higher transmembrane electrostatic potential. These patches lasted for ∼10 minutes, suggesting stable CHGB function. We also found that NaF and 0.5 mM CaCl_2_ in the *cis* side helped minimize leak currents. With 150 mM F^−^ or Cl^−^, the measured single channel conductance of CHGB is ∼125 pS (Fig. 3B).

**Figure 3.**
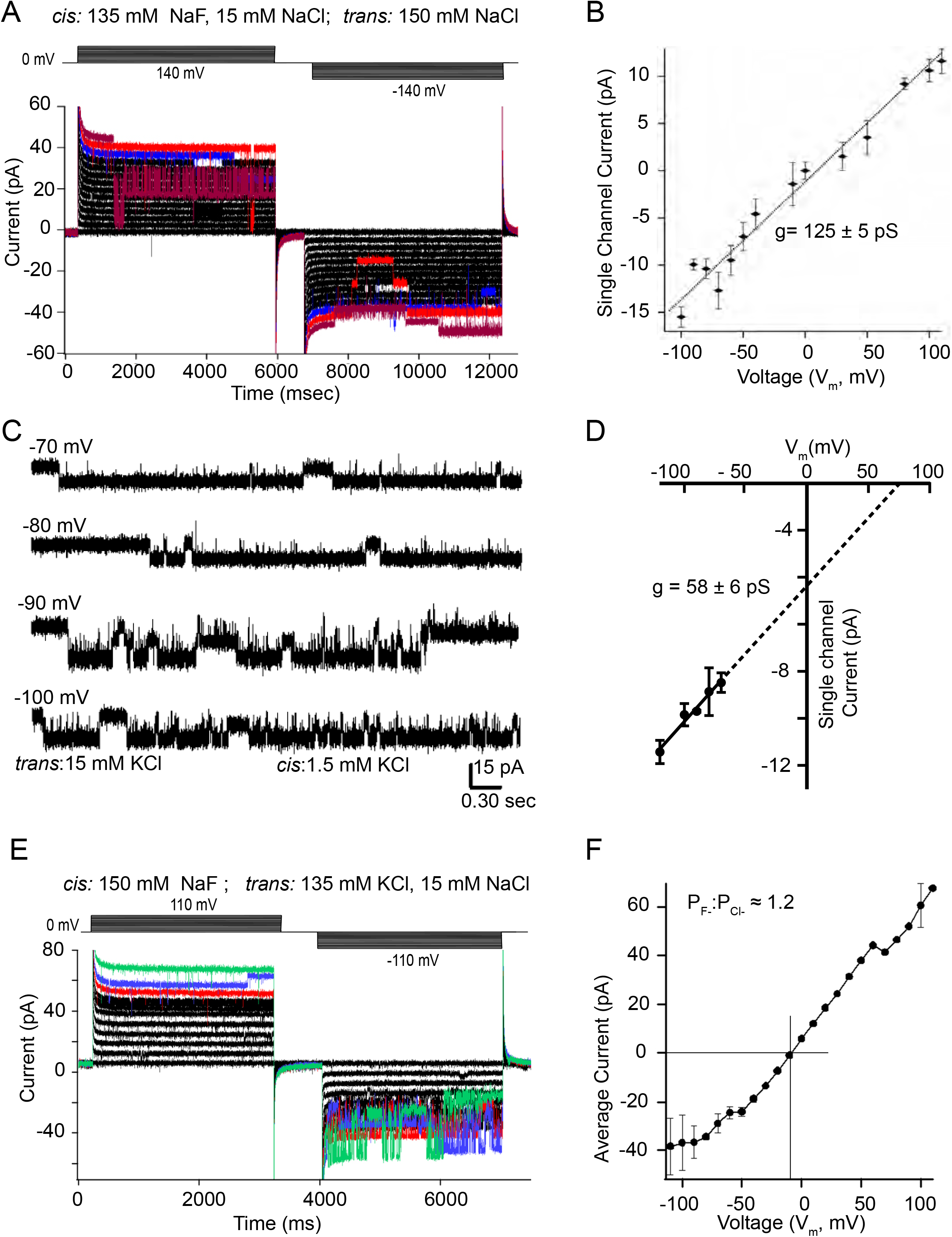
CHGB forms a chloride channel in membrane. **A)** Recordings from a multi-channel membrane in a planar lipid bilayer. Solutions were *cis*: 135 mM NaF, 15 mM NaCl, 0.5 mM CalCl_2_, 5.0 mM MES pH5.5; *trans:* 150 mM NaCl, 5.0 mM MES pH5.5. Holding potential: 0 mV; pulses in 10 mV steps to ±140 mV and ±140 mV in two epochs, respectively. Recordings at ±140, ±130, ±120 mV were colored dark brown, red and blue, respectively. (**B**) Single channel currents vs. transmembrane voltage (*V_m_*). Error bars: *s.d.* from 3-5 histogram-based measurements of single channel currents under each *Vm*. Fitting of the data with a linear function (dotted line) yielded a single channel conductance of 125 ± 6 pS with 150 mM F^−^/Cl^−^. (**C**) Single channel recordings from CHGB under low, asymmetric Cl^−^ and high, symmetric K^+^. *cis*: 150 mM K-isethionate, 1.5 mM KCl, 10 mM MES, pH 5.5; *trans*: 135 mM K-isethionate, 15 mM KCl, 10 mM MES, pH 5.5. (**D**) Single channel currents from (**C**) vs. *V_m_*. Without sizeable currents for *V_m_* varying from −50 to +100 mV under the solution conditions, a linear fitting (solid black line) was extrapolated to positive *V_m_* to yield estimated single channel conductance ∼58.4 pS and reversal potential ∼+65 mV (**E**) Small macroscopic currents recorded from a bilayer with multiple channels. The *cis* solution had 150 mM NaF, 0.50 mM CaCl_2_ and 5.0 mM MES pH5.5, and the t*rans* side 135 mM KCl, 15 mM NaCl, and 5.0 mM MES pH5.5. Holding potential 0 mV, and pulses in +10 and −10 mV steps (top). Recordings at ±110 mV, ±110 mv and ±90 mV were colored in green, blue and red to show the channel switching events. (**F**). Steady state currents from the recordings in (**E**) plotted against *V_m_*. Currents become smaller at negative range, due to more frequent channel closing (error bars are *s.d.*, n=4). The reversal potential (*E_rev_*=-8.0 mV) was used to estimate the permeation ratio (*P_F_ / P_Cl_*) of F^−^ : Cl^−^ using a simplified GHK equation, *E_rev_ = -RT/F ln(P_F_[F^−^]_o_/P_Cl_[Cl^−^]_i_*). The *cis* side is equivalent to the luminal side (outside) where the CHGB resides. From 8 measurements, *P_F_ / P_Cl_* =1.2± 0.2.

To test the Cl^−^ selectivity of the CHGB channel, we recorded single channel currents with asymmetrical Cl^−^ and symmetrical K^+^ at pH 5.5 (15 vs 1.5 mM KCL, Fig. 3C). K-isethionate was added to maintain high osmolality and ionic strength. Under such conditions, the recorded channels had no measurable outward current in the positive voltage range, suggesting no detectable conduction of K^+^. Analysis of the average single channel currents at different negative voltages yielded an estimated chord conductance of ∼58 pS (black line in Fig. 3D, for 15 mM Cl^−^) and by extrapolation a reversal potential of ∼+65 mV, close to the calculated Nernst potential (+59 mV) of Cl^−^, far away from that (0 mV) of K^+^, H^+^ or OH^−^. The detected channel is thus Cl^−^-selective with no K^+^ conduction. As negative controls, vesicles prepared with BSA, CHGA (Fig S4I), a CHGB deletion mutant lacking the CHGB-MIF (CHGBΔMIF) and CHGB-MIF all failed to generate any channel activity (data not shown), suggesting that the observed activity is probably genuine to CHGB. When we measured reversal potentials of the channels in bi-ionic conditions, the estimated permeation ratio between F^−^ and Cl^−^ (Fig. 3E, 3F) is ∼1.2. Because of the small number of channels in these recordings, we made sure that in higher voltage ranges, the channel closure reached zero current level briefly (as demonstrated by the red trace in Fig. S5A) such that the reversal potential determined from the current-voltage curves (Fig. 3F) was reliable for measuring the bi-ionic permeation ratio. Expanded traces (Figs 2C, S5A and S5B) showed subconductance states in different transmembrane potentials, which need further characterization in the future.

Because only a small number of channels were measured in lipid bilayers, we questioned if a trace amount of contaminating channels might have contributed to the recorded activities. We separated 20 micrograms of purified CHGB that had been kept in cold room for ∼4 days, and detected two faint smaller bands (1 and 2 in Fig. S5C). When the CHGB bands and the two shorter bands were cut out and digested for HPLC/MS analysis and Proteomic Identification (Supplementary Tables S1 and S2), all three bands were heavily dominated by CHGB peptides. The two shorter bands shared 5 peptides and were therefore CHGB degradation products. Other candidates had much fewer matched peptides, less than 6 for the CHGB band and only one for two smaller bands. Among them only KvQT member 5 (accession number E9Q9F in Table S1) is a K^+^ channel, but no anion channel. When we compared the scanned density of the bands (Fig. S5C, right), the two degradation bands, usually absent in fresh samples (Fig 1B), were ∼1.5% of total CHGB. Results from diluted proteins in the same gel suggested that any contaminant that is more than 0.12% of the total mass would have been detected. The purified CHGB thus has > 99.8% purity, making it very unlikely (<0.2%) for a contaminating anion channel to account for the observed channel activity in lipid bilayers.

### CHGB conducts F^−^ and Cl^−^ better than other anions

Because it was critically important to rule out stringently the possibility of trace contaminant channels yielding the measured channel activity in Fig. 3, we quantified anion flux from a large number (> 1.0E11) of CHGB vesicles. The Ag/AgCl measurement in Fig. 2C was limited to Cl^−^, not other anions; nor was it highly stable due to slow mixing of valinomycin and drifts of electrode potential. We instead implemented a light scattering-based flux assay (Fig. S3G) (Stockbridge et al., 2013) by both steady-state and stopped-flow fluorimetry. In both systems, valinomycin triggers K^+^ efflux and anion release from vesicles, resulting in a sudden drop of intravesicular osmolality and collapse of vesicles into ellipsoids that scatter more light (Fig. S3G, right side). Extrusion was used to control vesicle dimensions so that our measurements would not be dominated by large-sized vesicles. As a positive control, the bacterial EriC Cl^−^/H^+^ cotransporter in vesicles yielded a robust increase in light scattering (Fig. S4H) (Stockbridge et al., 2013). The CHGB vesicles prepared in parallel delivered a strong signal (Fig. 4A). The increase in steady-state light scattering saturated within the 10-s break after valinomycin addition, faster than the 20-s duration of Cl^−^ efflux in Fig. 2C. A non-specific Cl^−^ channel blocker, DIDS was able to block the strong signal with an apparent *K_d_* = 0.5 M (Fig. 4B).

**Figure 4.**
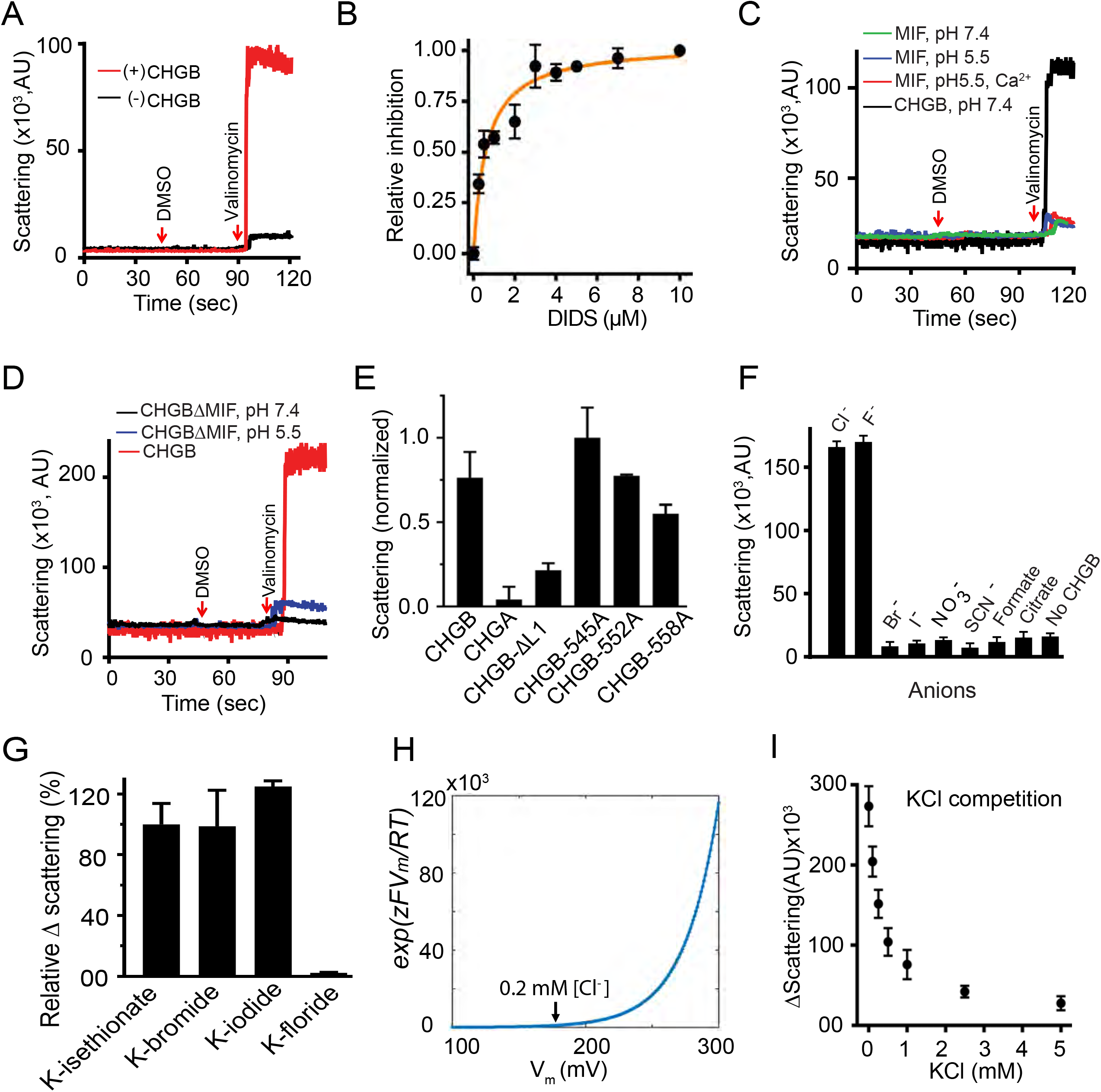
CHGB conducts F^−^ and Cl^−^ better than other anions. (A) Light-scattering signals measured from liposomes with (red) and without (black) CHGB at pH 7.4. A 30-sec pause (break in trace) for adding DMSO or valinomycin. (B) DIDS inhibits CHGB-mediated Cl-flux. Fitting with a Hill equation yielded a *k_d_* = 0.9 μM; n = 1.0. (**C**) Flux assay from liposomes with and without CHGB-MIF under different conditions in comparison with CHGB vesicles (black). (**D**) Flux assays of liposomes with and without CHGBΔMIF at two different pH’s. Wild-type control in red. (**E**) Flux assays of liposomes containing CHGA, CHGB and CHGB mutant proteins. Residues E545, E552 or E558 in the loop (L1) between helix2 and helix 3 was mutated to alanine. (**F**) Relative anion selectivity of CHGB channel based on the flux assay. Error bars represent *s.d.* from three different sets of experiments. (**G**). Instead of 300 mM K-isethionate, 300 mM KF, KBr or KI was introduced to the extravesicular side against 300mM KCl-loaded vesicles. Br^−^ or I^−^ supported significant flux signal. But 300 mM F^−^ abolished the flux. Error bars: *s.d.*, n = 3. (**H**). Calculated correction factor due to steady state Nernst potential for 300 mM KCl-loaded CHGB vesicles with 0-5 mM KCl outside. (**I**) Flux assay from CHGB vesicles in presence of 0-5 mM KCl in the extravesicular side with 300 mM K-isethionate and 1:10,000 PLR of CHGB: egg PC. Three independent sets of vesicles prepared in parallel (n=3) were separately measured.

As negative controls, recombinant CHGA in vesicles produced no signal (Fig. S4I-J); nor did CHGB-MIF or CHGBΔMIF (Fig. 4C-D). Hence, CHGB-MIF alone is insufficient to form the anion channel although it is indispensible to a functional channel. Considering the negatively charged residues contributing to anion selectivity of other Cl^−^ channels (Maduke et al., 2000), we deleted a short loop (residues 540-551; CHGB-ΔL1) right before Helix 3 (Fig. S4C) and mutated three negatively-charged residues in this loop to Ala (marked with arrows in Fig. S4C-D; Fig. 4E). CHGB-ΔL1 had significantly lower flux than wild-type CHGB. Mutation of the well-conserved E558 blocked approximately 50% of flux while E545A showed no effect and E552A had mild impairment (Fig. 4E). These data further demonstrate Helix 3 and the loop right ahead of it are important for ion conduction of the CHGB channel (Fig. S4D).

We next compared the relative ion selectivity of the CHGB channel by loading different anions into vesicles. Our data show that the CHGB channel conducts Cl^−^ and F^−^ much better than Br^−^, I^−^, NO_3_^−^, SCN^−^, formate or citrate (Fig. 4F), suggesting that the channel is much more selective than other known Cl^−^ channels or transporters (Fahlke, 2001; Maduke et al., 2000). Relative flux of F^−^ and Cl^−^ is consistent with the measured permeation ratio *P_F_/P_Cl_* =1.2 (Fig. 3F). An apparent physiological significance for higher anion selectivity of the CHGB channel is to prevent small intracellular organic anions (metabolites) from being passively concentrated into secretory granules and dumped as a waste.

The ion selectivity data (Fig. 4F) make two predictions. One is that extravesicular 300 mM Br^−^ or I^−^ should not affect the light-scattering signal, while F^−^ should completely stop it, precisely what our experiments showed (Fig. 4G). The other is that a low concentration of extravesicular Cl^−^ should shift the initial steady-state Nernst potential across vesicle membranes and significantly inhibit Cl^−^ flux because of a voltage-dependent factor affecting the rate of valinomycin-mediated potassium flux (Fig. 4H; Section 18 in Supplementary information). Indeed, extravesicular [Cl^−^] of 0.1 to 2 mM significantly impaired the light-scattering signal (Fig. 4I).

### High-cooperativity among CHGB subunits in forming functional channels

Steady-state fluorimetry measurement cannot detect fast channel-opening events because of slow and uneven equilibration of valinomycin among vesicles. A stopped-flow fluorometer (Hayner et al., 2014) with a dead time of 2 ms was used to overcome this limit. We compared the light-scattering signals by both steady-state and stopped-flow fluorimetry, and titrated PLRs of CHGB vesicles with ∼0.4 mg/ml lipids. When PLR < 1:50,000, no signal was detected in either system (Fig. 5A-B). When PLR reached 1:1,000, the signal approached maximum. All stopped-flow traces showed instantaneous jump, suggesting a fast efflux of K^+^/Cl^−^ within 2 ms. Because of fast and even partitioning of valinomycin into vesicles, the instantaneous jump in light scattering reflected a sudden change in vesicle shape after KCl efflux. When measurements from the first 5 seconds from the steady-state fluorimetry and the first 40 ms of the stopped-flow data (Fig. 5C & 5D) were plotted against [CHGB] and fitted with a Hill equation (Fig. 5E), the estimated Hill coefficients were ∼1.2 from the steady-state measurement and ∼4.2 from the stopped-flow data, suggesting high positive cooperativity among CHGB subunits in forming a functional channel. The steady-state data revealed less cooperativity probably because of uneven and slow mixing of valinomycin with vesicles and the complication from water movement in response to osmolality change. When the data in Fig. 5E were plotted against the average number of CHGB subunits per 100 nm vesicle (Fig. 5F), the threshold for strong signals occurred at ∼4 CHGB subunits per vesicle, and nearly 80% of the maximum signal was achieved with ∼8 CHGB subunits per vesicle. Assuming a Poisson distribution of CHGB subunits in vesicles, our data suggest that most probably 4 CHGB subunits form one channel (Section 19 in Supplementary Information).

**Figure 5.**
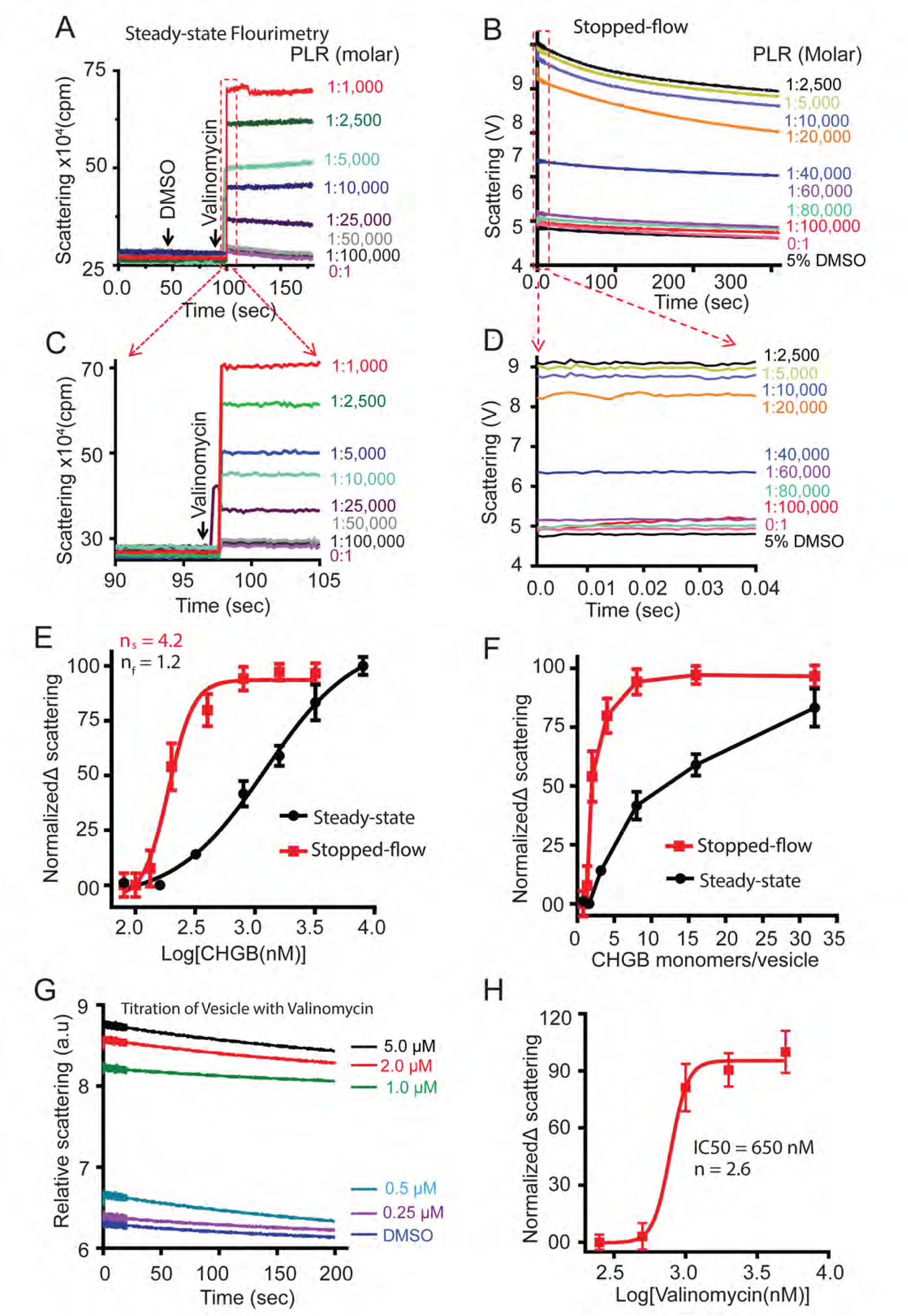
Fast kinetics of Cl^−^ release from the CHGB channels in vesicles. (**A**) 300 mM-KCl-loaded vesicles changed into 300 mM K-isethionate right before the assay. Time-lapsed light scattering measurement at 600 nm in a Fluormax-4. Vesicles of different PLRs were prepared in the same batch. Adding DMSO triggered no signal. Adding 1.0 μM valinomycin does. (**B**) Time-lapsed light scattering (610 nm) measured in a stopped-flow system, by mixing equal volumes of vesicles and 2.0 μM valinomycin in buffer. 5% DMSO solution was added as negative control. Vesicles of PLR=1:1,000 used to set a proper accelerating voltage for the PMT tube to have the broadest dynamic range. (**C**) Expanded region in the red box in panel **A**. (**D**) Expanded region in the red box in panel **B**. Data from the first 40 ms after the dead time of ∼2 ms for mixing are shown. (**E**) Average signals from the first 5-second data in **C** (black) and those from the first 40-ms in **D** (red) are plotted against [CHGB]. Fitting with Hill equations (solid lines) generated different Hill coefficients. Steady state data from four different datasets (n=4; error bars are *s.d*.); Stopped-flow data from two sets of triplicate measurements (n=6; error bars are *s.e.m*.). (**F**) The data in panel E were plotted against the average number of CHGB monomers per 100-nm vesicle. The threshold for generating a significant signal is ∼4 monomers per vesicle for both measurements. (**G**) Titrating valinomycin for the light-scattering from CHGB vesicles in the stopped-flow system. The CHGB vesicles had PLR = 1:25,000. More vesicles than those in Panel B were used to boost the signal, which also increased the baseline signal (Buffer_DMSO). Three triplicate measurements were obtained (n=9; error bars are *s.e.m.*). (**H**) The normalized signal from panel G were plotted again [Valinomycin] and fitted with a Hill equation to yield a Hill coefficient of ∼2.6.

Valinomycin might form a K^+^ channel. The light-scattering assay requires fast K^+^ flux. Varying [valinomycin] per vesicle is hence expected to affect the signal. Titration of [valinomycin] in the stopped-flow measurement (Fig. 5G) found that when [valinomycin] <0.25 μM, the flux signal was almost non-detectable, whereas 2-5 μM delivered the maximal signal. Fitting of the dose response data with a Hill equation (Fig. 5H) identified a Hill-coefficient of *n*=2.6, indicating high cooperativity among valinomycin molecules in moving K^+^ across membrane. These data accord with our observation that 0.1 μM valinomycin triggered no detectable flux signal. The cooperativity suggests that valinomycin function as trimers or high-order oligomers in membrane, reminiscent of the dimeric valinomycin channel in ultrathin membranes (Gliozzi et al., 1996). To match the Cl^−^ flux, each valinomycin oligomer must have a conductance > 3 pS, a lower limit for an uncharacterized valinomycin channel.

### CHGB channel is required for granular maturation in neuroendocrine INS-1 cells

Because CHGB is a housekeeping protein in secretory granules (Brunner et al., 2007), we tested whether its channel can serve the anion conductance first recognized in chromaffin granules as essential for granule acidification(Johnson et al., 1982). Because the secretory granules are too small (mostly <500 nm) for direct patch-clamp recording, no specific inhibitor for the CHGB channel is available yet, and prior attempted recordings from purified granules all suffered from certain level of contamination by other cell membranes (Guzman et al., 2013; Inagaki et al., 2010; Kelly et al., 2005), we instead used genetic manipulations to alter CHGB, a ratiometric fluorophore to measure intragranular pH (Stiernet et al., 2006) and (semi-)quantified proinsulin and insulin in INS-1 cells.

To perform ratiometric pH measurements, we first examined whether a Lysosensor DND-160 preferentially stained secretory granules in INS-1 cells. A fluorescently labeled secretory granule protein, syncollin-pHluorin (Fernandez et al., 2011), was expressed (Fig. S6A) and imaged together with DND-160-stained acidic compartments in live cells. DND-160 overlapped very well with syncollin, meaning that nearly all DND-160-stained compartments were secretory granules, which is consistent with published observations(Stiernet et al., 2006).

To alter the CHGB proteins, siRNAs were used to suppress the endogenous CHGB, which allowed transient overexpression of CHGB or its mutants that supersede the siRNA effects. We found that 100 nM CHGB siRNAs (Zhang et al., 2014) (Fig. S6B) suppressed >98% of CHGB protein (Fig. S6C). To evaluate possible OFF-target effects, four individual CHGB siRNAs in the mixture were tested separately. All four were effective with siRNA 2 & 4 being ∼2-fold more effective (Fig. S6D). We therefore used these two siRNAs individually or in combination to minimize OFF-target effect.

To measure intragranular pH, both dual-excitation (Fig. S6E) and dual-emission experiments (Fig. 6A) were performed. CHGB knockdown reduced significantly the total number of secretory granules per cell (the second row in Fig. 6A and the bottom row in Fig. S6E). Control siRNAs with scrambled sequences (CTL siRNAs) had no effect (top row in Figs 6A and S6E). Cells treated with NH_4_Cl were used as a positive control because NH_4_^+^ increases intragranular pH to near neutral or higher. 100-140 granules were randomly selected from paired images of multiple cells to measure ratios in fluorescence (Section 16 in supplementary information). Histograms of the ratiometric measurements show a clear shift in pH distribution (blue vs. grey bars in Fig. S6F). Histogram comparison is thus a robust and objective strategy. Comparison of average granular pH values in differentially treated cells (Fig. S6G) showed that the CTL siRNAs caused no significant change in intragranular pH whereas CHGB knockdown caused significant deacidification in the leftover secretory granules (red and grey in Figs S6F & S6G). The same results were obtained from dual-emission experiments (485 nm and 510nm, top two rows in Fig 6A, black vs. red in Fig. 6B and histograms in Fig. S7A). We continued with dual emission experiments to avoid the need of a quartz objective lens for near-UV excitation.

**Figure 6.**
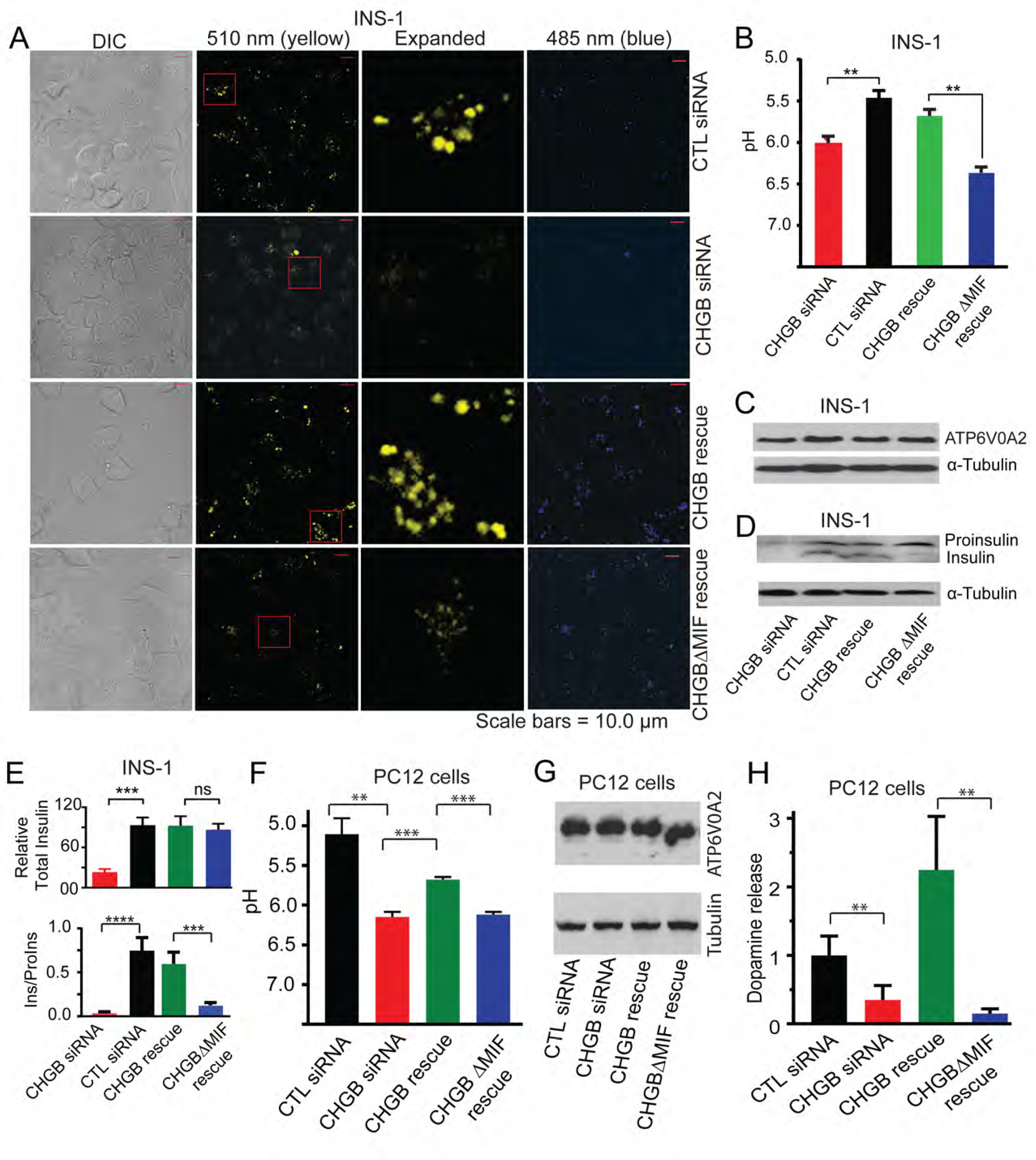
CHGB channel required for granule acidification and insulin maturation in INS-1 cells. (**A**) Ratiometric pH measurements. INS-1 cells on a glass-bottom dish (DIC), transfected with control siRNAs (top row) or CHGB siRNAs (rows 2 to 4). For the top two rows, cells were stained after 96 hours with one medium change after 48 hours. For rows 3 & 4, 48 hours after transfection with CHGB siRNAs, cells were transfected with plasmids carrying CHGB or CHGBΔMIF and imaged after 48 more hours. Excitation: 410 nm; emission: 485 nm and 510 nm. A small area (red window) in each 510 nm image expanded to show the granules. Measurement at 510-nm in row 4 was weaker due to pH difference. (**B**) Average pH from hundreds of secretory granules from cells described in **A**. (**C**) Western blot for ATP6V0A2 (vesicular H^+^-ATPases) from cells as in (**A**). (**D**) Western blot for both insulin and proinsulin from lysates of cells as in (**A**). (**E**) Relative total insulin (insulin + proinsulin; Upper panel) and insulin maturation (Ins / ProIns; lower panel) from cells as in (**A**). Relative total insulin was normalized to cells treated with control siRNAs (black). Error bars are *s.d.* in **B** (n > 100, Fig. S6A & B) and **E** (n=3). **: *p* < 0.05; ***: *p* < 0.001; ****: *p* < 0.0001; *ns*: not significant. (**F**) Average pH from > 100 randomly selected granules (Histograms in Fig S7C & D). Error bars: *s.d.* (n>100). (**G**) Western blots for ATP6V0A2 in four differentially treated PC12 cells. (**H**) Relative dopamine content in the granules released from depolarization-treated cells. An ELISA kit was used. Standard *t*-test for F and two-tailed Welch’s *t*-test for G. **: *p* < 0.05; ***: *p* < 0.001.

Next we compared wild-type CHGB and nonfunctional CHGBΔMIF by transiently overexpressing them in CHGB-knockdown cells (Fig. 6A, rows 3 & 4). Both constructs had the N-terminal Cys-loop and could support granule biogenesis, certifying their overexpression and their mRNAs outcompeted the siRNAs. Comparison of intragranular pH’s (Fig. 6B, green vs. blue; Fig. S7B) found that wild-type CHGB restored granular acidification, close to cells treated with CTL siRNAs (black, Fig 6B). Contrastingly, CHGBΔMIF failed to restore granule acidification (blue in Fig. 6B & Fig. S7B). Because CHGBΔMIF supported granule biogenesis (Fig. 6A), but had no channel function (Fig. 4D), these findings demonstrated directly that granule acidification in live cells needs CHGB channel function.

De-acidification of secretory granules might alternatively result from decreased activity of vesicular H^+^-ATPase. To test this possibility, we examined the expression of a key vATPase subunit, ATP6V0A2, and found no change among the four differentially treated cells (Fig. 6C). This result was expected because our CHGB siRNAs were on target (Fig. S6D) and did not affect the vesicular H^+^-ATPase.

Granule deacidification is expected to slow down insulin maturation (Davidson et al., 1988). When proinsulin and insulin were detected by western blot from the four differentially treated cells (Fig. 6D, E). CHGB knockdown decreased total insulin (Ins + ProIns; top in Fig. 6E) and impaired insulin maturation (Ins / ProIns; bottom in Fig. 6E). Overexpressed CHGB corrected both defects. But CHGBΔMIF overexpression restored only total insulin, not insulin maturation (Fig. 6E) because it lacked channel function and failed to acidify the granules for normal insulin maturation.

### CHGB channel affects dopamine loading in PC-12 cells

To evaluate whether the CHGB conductance is generic to endocrine cells, we tested it in PC-12 cells, which contain the same machineries for regulated secretion as adrenal chromaffin cells. Intragranular pH’s among four differentially-treated PC-12 cells were compared (Figs. 6F, S7C, S7D). Similar to findings in INS-1 cells, CHGB knockdown decreased the number of secretory granules significantly and overexpressed CHGB or CHGBΔMIF restored it. CHGB knockdown de-acidified the granules. Overexpression of CHGB reversed it, but overexpressed CHGBΔMIF failed to do so (blue vs. green bars in Fig. 6F). The genetic manipulations did not change ATP6V0A2 expression (Fig 6G). The CHGB channel function is thus a must for normal granule acidification in PC-12 cells, too.

Concentrating dopamine in PC-12 cells requires CHGB channel. H^+^-coupled vesicular monoamine transporters pump dopamine into secretory granules (Wimalasena, 2011). Granule deacidification was expected to decrease intragranular concentration of dopamine. We measured dopamine content in readily releasable granules (RRGs) from depolarization-stimulated PC-12 cells using an ELISA kit (Fig. 6H) (Nishimura et al., 2010). Statistical comparison showed that CHGB knockdown significantly decreased dopamine content in RRGs, which was recovered by CHGB, but not CHGBΔMIF (blue vs. green bars in Fig. 6H). The CHGB channel in PC-12 cells is therefore required for proper loading of dopamine into secretory granules.

### CHGB remains on cell surface after exocytotic release of insulin-granules

Because all known CHGB extracellular functions are mediated by its processed peptides, we questioned whether the full-length CHGB protein was retained after granule release. In PC-12 cells, the “tightly membrane-associated” CHGB stayed on cell surface for 2-4 hours and recycled back into the cells(Pimplikar and Huttner, 1992). We tested if the same is true for INS-1. After INS-1 cells were treated with high K^+^, a small fraction (<0.5%) of granules were released. CHGB-antibodies were used to label CHGB on the surface of the non-permeabilized cells at 0-4 degree C, which stopped endocytosis. Alexa 488-labeled secondary antibodies were used for visualization under a confocal fluorescence microscope (FM). Multiple puncta of CHGB showed on the surface of the positive cells (Fig. 7A), suggesting that after exocytosis, full-length CHGB proteins remain membrane-integrated at the exocytotic sites. Quantification of surface puncta per cell shows that the positive cells have on average 9-10 CHGB puncta, significantly higher than the control cell (2-3; Fig. 7B-C). Similarly, when INS-1 cells were treated with glucose, specific labeling with fluorescent antibodies yielded much stronger signal on the labeled than control cells (Fig. S7E). The surface retention of full-length CHGB is a good way to separate it from its processed peptide hormones.

**Figure 7.**
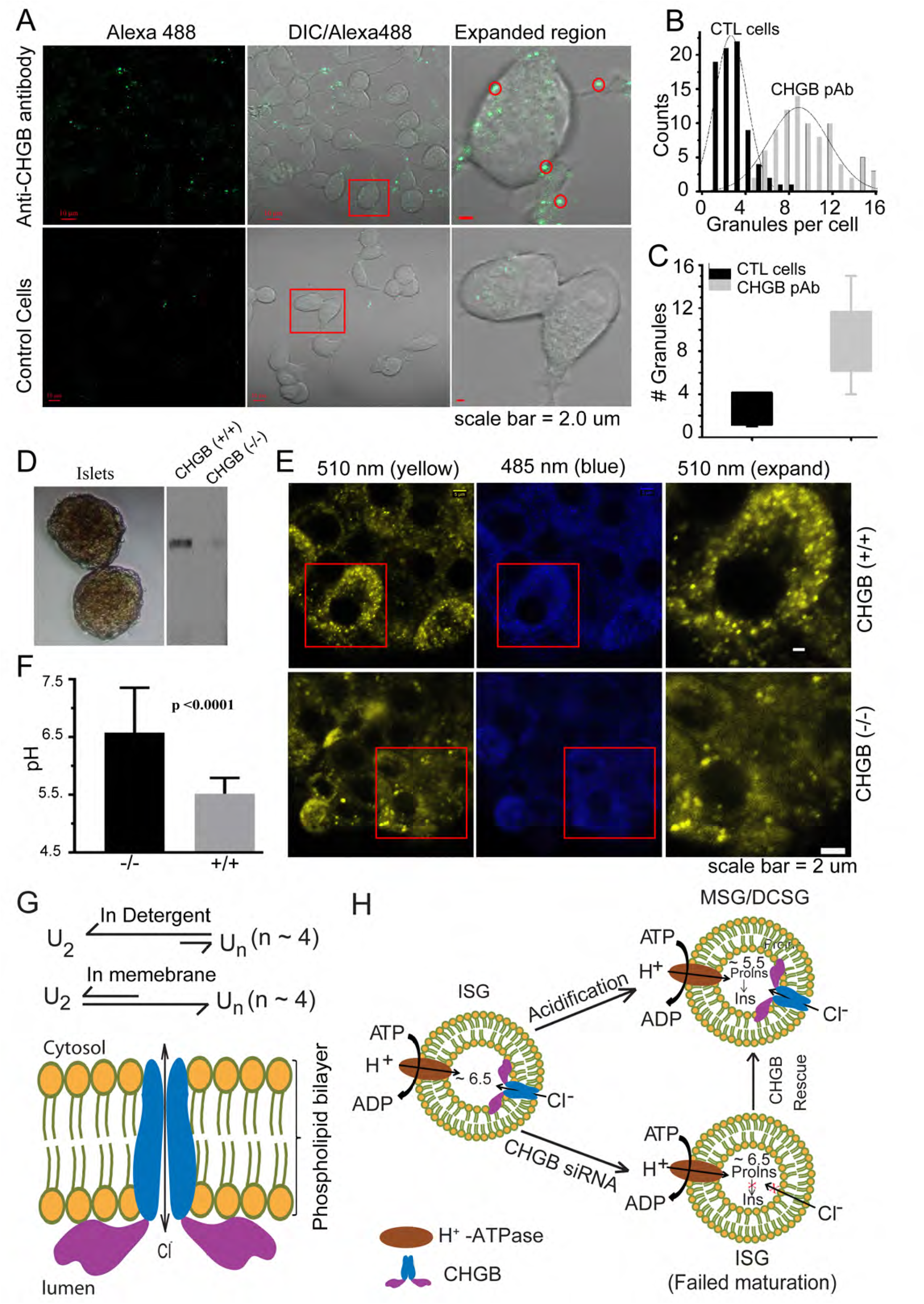
A working model for CHGB membrane insertion and its anion channel function in regulated secretion. (**A**) INS-1 cells incubated with high KCl (55 mM) for 15 min at 37ºC to release granules before being labeled with anti-CHGB antibody on ice. Alexa 488-conjugated secondary antibody was used for visualization of CHGB puncta at the live cell surface by confocal immuno-FM (63x objective). Upper row shows images from positive cells. Lower panel from cells treated only with 2^nd^-Ab. (B) CHGB punta per cell are compared between two conditions (# of cells >50). (C) Average number of Alexa-488 punta for the two conditions. (D) Two typical pancreatic islets (left) and western blot of CHGB in liver tissue from wild-type (CHGB+/+) or from knockout mouse (CHGB−/−). (E) Ratiometric pH measurements in islet beta-cells of CHGB+/+ vs. CHGH−/− mice. 10 islets of each genotype were stained with DND 160 and imaged under Zeiss LSM-800 confocal microscope (63x objective). Excitation: 410 nm; emission: 485 nm and 510 nm. A small area (red window) in each 510 nm image expanded to show the granules. Beta-cells from knockout mice have fewer granules and obviously fainter signal in the 510 nm channel due to granule deacidification. (**F**) Average intragranular pH in mouse beta cells of two different genotypes (CHGB−/− vs. +/+). (**G**) Dynamic equilibrium exists between CHGB dimers and oligomers (probably tetramers, U_n_). In detergents dimers are favored while in membrane oligomers are. Bottom: a scheme for CHGB insertion into membranes from the luminal side, forming a Cl^−^ channel and part of each CHGB subunit resides on the luminal surface. (**H**) CHGB channel, together with the vesicular H^+^-ATPase, supports normal acidification of secretory granules. CHGB knockdown impairs acidification and slows down cargo maturation, and overexpressing CHGB restores it. Insulin maturation is shown as an example for cargo maturation.

### Granule acidification in pancreatic beta-cells in CHGB knockout mice is impaired

If the mechanism for CHGB channel is native to secretory granules, granule acidification should be affected in the CHGB knockout mice. We raised a knockout mouse strain produced by the Wellcome Trust Sanger Institute and distributed by EMMA (#10088; URL: https://www.infrafrontier.eu/). Wild-type and homozygous littermates were raised for experiments. Female and male mice were analyzed separately in parallel. Fig. 7D showed the isolated islets and the detection of CHGB from liver tissues of wild-type (CHGB+/+) and knockout (CHGB−/−) mice. Genotype confirmation was done for every mouse (Fig. S8A). Western blot detection was performed for different types of tissues in order to confirm the penetrance of the knockout mutation. Freshly isolated islets were stained with DND-160 and imaged. To avoid possible variation in staining, we focused on the cells closer to the surface of each islet. The wild-type beta-cells (top row of Fig. 7E) have much more well-stained granules than the cells from the knockout mice (bottom row of Fig. 7E; contrast of these images was enhanced to show the otherwise faintly-stained granules with increased background; raw data were used for ratiometric measurements). Ratiometric measurements from hundreds of granules found that CHGB knockout caused significant deacidification in the insulin-secretory granules in pancreatic beta-cells from both male and female mice (Fig. 7F; S8B).

## DISCUSSIONS

### CHGB-membrane interaction and channel activity in secretory granules

Our data collectively support a working model that CHGB inserts itself into lipid membranes from the luminal side, which promotes CHGB oligomerization by shifting a dynamic equilibrium between the dimers and the predicted tetramers, and generates a Cl^−^ channel (Fig. 7G). Because cells have sufficient ER membranes, nascent CHGB molecules should be all membrane-associated from the luminal side. Both helices 1 and 3 are involved in the membrane insertion (Figs 2I-J) and the loop before helix 3 is important for anion conduction (Fig. 4E). Moreover, the CHGB-membrane interaction may promote high-order oligomeric forms in a calcium-dependent way (Fig. S2D). For simplicity, we depict a channel pore in a dimeric fashion, omitting other subunits in Fig. 7G. Once the channel is formed, it opens most of time, making the granular membranes highly permeable to Cl^−^, but not other intracellular anions. If the CHGB channel function is abolished, lack of Cl^−^ influx into the granules impairs granule acidification and slows down the conversion of proinsulin to insulin in INS-1 cells or pancreatic beta-cells (Fig. 7H) or decreases the H^+^-coupled pumping of dopamine in PC-12 cells. Overexpressed CHGB restores granule acidification and maturation, but its nonconducting deletion mutant does not.

In accord with our model and the impaired granule acidification in the CHGB knockout mice, hyperproinsulinemia and insulin secretion defects were reported in a different knockout mice (Obermuller et al., 2010). Similarly, in another knockout mice a significant decrease of catecholamine content was reported in released chromaffin granules (Diaz-Vera et al., 2010), and was attributed to the saturation of an unknown intracellular mechanism for dopamine concentration. Our model now depicts a concrete mechanism that without CHGB the chromaffin granules de-acidify and pump much less dopamine from cytosol to granules.

### Membrane-insertion induces CHGB channel formation

In neuroendocrine cells, CHGB is co-translationally transferred into ER, where it should be inserted in membrane. Similarly, our data suggest that CHGA is fully membrane-attached (Fig. S4I). Our data show that both CHGB and CHGA are fully reconstituted in the liposomes so we propose that the full-length CHGB is always in membrane in the regulated secretory pathway. Because both ER and Golgi membranes have their own Cl^−^ channels, the CHGB channel is nonessential for ER and Golgi. The Cys-loop domain of the CHGB guides its delivery to its indispensible role in the secretory granules. Further, partial processing of a fraction of CHGB proteins into short peptides in secretory granules should render a significant number of anion channels defective before granule release. Without studying whether the “tightly membrane-associated” CHGB on cell surface is functional (Pimplikar and Huttner, 1992), we propose that CHGB channel’s main function is intragranular and may be tightly regulated. After granule release, the full-length CHGB is retained in membrane and is recycled back into the granules, which separates the intracellular functions of the CHGB channel from the extracellular functions of CHGB peptides, especially those that act as peptide hormones.

Membrane insertion of a protein without canonical hydrophobic transmembrane segments to form ion channels has multiple precedents. Hemolysin, C-type lectin, VopQ, etc. are a few (Mukherjee et al., 2014; Ramarao and Sanchis, 2013; Sreelatha et al., 2013). Thermodynamically, oligomerization of individual subunits and interactions among them and with lipids increase entropy of water molecules liberated from hydrophobic surfaces, such as those from helices 1 and 3 in CHGB (Fig. 2I-J), and overcome energetic cost of dehydration and membrane insertion. CHGB-coated 25 nm nanoparticles may reconcile contradictory results between native CHGB being partially purified as soluble (heat-stable) fractions and its tight interaction with lipid membranes (Benedum et al., 1986; Benedum et al., 1987; Pimplikar and Huttner, 1992; Yoo, 1995b). The CHGB-packed 25nm nanoparticles are expectably heat-stable and soluble by burying CHGB’s hydrophobic domains. It would be interesting to examine whether CHGB in the soluble (or heat-stable) fractions takes similar structures as in the 25 nm nanoparticles (Fig. S2B-C). Similarly, structures of CHGB dimer, tetramer, and higher-order oligomers will be needed to reveal their allosteric changes during channel formation. Further, the sidedness of CHGB insertion in membrane endows special functions such as membrane remodeling and pH-or Ca^2+^-dependent regulation, which await more investigations.

A “tightly membrane-associated form” of the CHGB is resistant to treatment of basic pH, but soluble by Triton X-114 (Pimplikar and Huttner, 1992). Beside a good mechanism to retain the full-length protein and separate it from the CHGB peptides, such a mechanism remains consistent with our working model because calcium-induced CHGB aggregates (Fig. S1C-F) or CHGB-packed nanoparticles might send some full-length proteins into the luminal side through membrane remodeling. It remains unknown what physiological roles the 25 nm particles, if they form inside the cells, would play. Lipid composition of the granular membranes may be another factor underlying CHGB-membrane interactions. Nevertheless, the CHGB channel activity is a generic property because of the channel activity in all membrane systems tested so far and its necessity to normal granule maturation in both cultured cells and primary pancreatic beta-cells. Considering the highly conserved helices (Fig. S4B, S4D) and the key residues (e.g. E558 in Fig. 4E, S4D), we propose that the Cl^−^ conduction represents a universal property within the CHGB subfamily in regulated secretory granules.

Our observations in neuroendocrine cells and primary beta-cells explain the main phenotypes of the CHGB knockout mice — hyperproinsulinemia and decreased catecholamine content in chromaffin granules (Diaz-Vera et al., 2010; Obermuller et al., 2010; Zhang et al., 2014). Future studies need to examine whether impaired CHGB channel contributes to defective insulin secretion in Type 2 diabetes and other neuroendocrine diseases because CHGB dysfunction is associated with hypertension, diabetes, Parkinson’s diseases, schizophrenia and Alzheimer’s diseases(Zhang et al., 2014) and its overexpression with multiple forms of endocrine cancers. Verification of a causative relation between altered CHGB-channel function and these diseases will make the CHGB channel a druggable target in the future.

### Separation of CHGB function in granule biogenesis, maturation and release

Without firmly verified physiological functions, CHGB was proposed to be involved in biogenesis, maturation and release of secretory granules. Its N-terminal Cys-loop is a sorting signal (Chanat et al., 1993), even though its sorting mechanism remains hypothetical (Hosaka and Watanabe, 2010). We observed CHGB-packed nanotubules and nanospheres (Figs 1F, S2B & S2C). It is tempting to speculate that the concentrated CHGB and other proteins at the TGN sorting sites could induce CHGB-packed nanostructures and might somewhat facilitate granule biogenesis through interacting with the luminal surfaces.

The CHGB anion channel serves an unprecedented role in granule maturation. Without the N-terminal Cys-loop sequence, the membrane-interacting part of CHGB might still form a channel, but would be mis-located in cells. CHGB therefore combines a strong sorting signal with a channel-forming portion in order to achieve proper delivery of the latter. The ClC3 transporter, being absent from the granules(Maritzen et al., 2008) and contributing no significant outward anion flow into the luminal (extracellular) side(Guzman et al., 2014; Guzman et al., 2013), may still affect proton-pumping or chloride flux through physical interactions and generate the phenotypes observed in the ClC3-null mice (Deriy et al., 2009; Li et al., 2009). Both ClC3-null and CHGB-null mice exhibited hyperproinsulinemia and lower catecholamine content in secretory granules (Diaz-Vera et al., 2010; Obermuller et al., 2010; Zhang et al., 2014). Whether the two are functionally coupled remains a mystery.

CHGB has been proposed to directly interact with the IP_3_Rs in the secretory granules and regulate granule release (Yoo et al., 2001; Yoo et al., 2000). It is possible that the CHGB Cl^−^ conductance may facilitate the fast release of Ca^2+^ from the granules before exocytosis. This role may be fulfilled by the outwardly rectified ClC-3 channel, should ClC-3 is delivered to the granule membranes. Further study will be needed to understand the possible relationship between CHGB and ClC-3.

In the CHGB knockout mice, exocytotic release of RRGs is normal because the CHGB channel is not required for vesicle fusion. ClC-3 is proposed to play a critical role in granule release (Deriy et al., 2009; Li et al., 2009). There are likely two possible roles for ClC-3 here: either it is required for facilitating Ca^2+^-efflux from the granules or it functions at the cell surface to facilitate Ca^2+^-influx before granular exocytosis. It is unclear whether CHGB contributes to either or both of these functions. However, it is equally possible that the intracellular ClC-3 or a plasma membrane ClC channel may be a licensing factor for exocytotic release of the secretory granules.

It is unclear whether the CHGB channel is important for its endocytotic uptake (Pimplikar and Huttner, 1992). ClC-3 is proposed to have dual functions, either as a Cl^−^/H^+^ exchange or as a Cl^−^ channel. It was found to be in early and late endosomes and in synaptic vesicles (Guzman et al., 2014). ClC-5 is detected in endocytotic vesicles, especially in the kidney proximal tubules (Gunther et al., 1998), suggesting that ClC-3 and ClC-5 might play similar roles in different cell types. When CHGB on the cell surface is endocytosed, its channel function would co-exist in the endosomes with ClC-3 or ClC-5(Pimplikar and Huttner, 1992). How the intracellular machinery separates these channels when CHGB is recycled back to Golgi membranes or secretory granules is an open question.

### Relations between CHGB and other proposed granular channels

Besides IP_3_Rs in secretory granules, there have been at least three distinct views on ion channels in the secretory granules. A large-conductance K^+^-channel (BK) and a few other K channels and a 250 pS Cl^−^ channel were detected in isolated chromaffin granules, in conflict with the early report of no cation permeability for such granules (Hordejuk et al., 2006; Johnson et al., 1982; Johnson and Scarpa, 1976). Kir6.1, ATP-sensitive K^+^/Cl^−^ channels and ClC1/2 channels were reported from zymogen granules, but these ClC channels showed completely different properties from ClC channels relocated to cell surfaces (Guzman et al., 2013; Kelly et al., 2005; Thevenod et al., 2000). All these conflicting results may be partially due to difficulty in avoiding contaminating intracellular or plasma membranes. Only in a well-controlled clean system can high-level certainty be achieved in order to resolve these controversies. Patch-clamp methods do not allow direct recordings from secretory granules due to their small size (mostly < 400 nm). We will need highly specific CHGB channel blockers in order to separate the CHGB channels from other Cl^−^ channels in cell membranes. Needless to say, specific CHGB channel inhibitors will have high clinical value when the connections between altered CHGB function and human diseases are fully solidified.

To conclude, our data for multiple angles converge to the conclusion that CHGB in membrane forms a chloride channel that is essential to normal granule acidification and cargo maturation in both cultured endocrine cells and primary beta-cells. Two conserved amphipathic helical segments mediate the membrane-induced channel formation. The channel function probably is a generic property for the CHGB subfamily. The anion selectivity of the CHGB channel is high and unique within this subfamily and its structural basis awaits further investigations.

## Methods

Details available in the supplemental information. *CHGB preparation and reconstitution:* Recombinant CHGB were purified and inserted into lipid vesicles by removing detergents with BioBeads.

*EM examination of CHGB-containing vesicles:* Glow-discharged carbon-coated grids were used to present CHGB vesicles for negative-staining and for imaging in an electron microscope.

*Single particle EM of CHGB dimers:* Individual dimers were images by negative stain and cryo-EM. Datasets of individual molecules were built for image analysis and 3D reconstruction. *Recordings of CHGB activities by three different methods:* CHGB fused in bilayers were recorded by patch-clamp methods. Cl^−^ releases from CHGB in vesicles were recorded with a Ag/AgCl electrode and in a light-scattering assay, respectively.

*Granule acidification and cargo maturation in live cells*: Cells were manipulated differently and intragranular pH’s were monitored by ratiometric measurements from DND-160 stained granules. Insulin maturation in INS-1 cells was monitored by Western blot. Dopamine content in granules released from the same number of PC-12 cells was measured by ELISA.

## Acknowledgements

Detailed acknowledgements are in supplemental information.

## Author Contributions

Q.-X.J. designed and oversaw the experimental studies, analyzed the results with all co-authors. Q.Y. did the initial molecular cloning of the CHGB cDNA and tested the expression of the CHGB protein in culture cells and in *sf9* cells, G.Y. developed most of the molecular biology and performed all biochemistry and cell based experiments, LGD and LBB guided the use of the stopped-flow system for the flux assay, and H.J. and Q-X.J. conducted all electrophysiological experiments. M.A. performed the dissection of mouse pancreatic islets. All authors contributed to data analysis and manuscript writing.

## Acknowledgements

We are grateful to Dr. Barbara Ehrlich (Yale University) for providing the constructs for CHGB, to Dr. Christopher Miller (Brandeis University) for sharing the construct for *EriC* and the protocols for the chloride flux assays, to Dr. Herbert Y Gaisano (University of Toronto) for the syncollin-pHluorin construct, to Dr. Kuixing Zhang (UC San Diego) for sharing his techniques for releasing granular contents from endocrine cells, to Dr. Sohini Mukherjee and Lora Hooper (UT Southwestern Medical Center) for letting us use their FluoroMax-3 system for studying the release of fluorophores from reconstituted vesicles, to Dr. Sandra Schmid (UT Southwestern Medical Center) for accessing a Horiba fluorometer, to Drs. Ilya Bezprozvanny, Kate Phelps, and Jen Liou for assistance and advice on the intragranular pH measurement using ratiometric dyes, to Dr. Wen-Hong Li for sharing the INS-1 cells and the insulin detection methods, and to Dr. Jerry Shay for providing the Lentiviral construct of CHGB shRNA. We thank Drs. Jose Rizo-Rey, Paul Blount, Peter Michaely and Lily Huang at UT Southwestern, Frederick Sigworth at Yale University, Drs. Zhonglin Mou and Julie Furlow-Maupin at University of Florida, and Michael X. Zhu at UT Health Science Center at Houston for critically reading and commenting on the manuscripts, and Dr. Christopher Miller for insightful discussions on the data and the interpretation of the results. We thank Drs Jin Koh and Sixue Chen in the Proteomics Core at the Interdisciplinary Center for Biotechnology Research (ICBR) of University of Florida for technical support in Mass Spectrometry and Proteomic analysis. The work was mainly supported by NIH (R01GM111367 & R01GM093271 to Q.-X.J.), CF Foundation (JIANG15G0 to Q-X.J), Welch Foundation (I-1684 to Q.-X.J.) and CPRIT (RP120474 to Q.-X.J.), and partially by an AHA National Innovative Award (12IRG9400019 to Q.-X.J.), an NIGMS EUREKA Award (R01GM088745 to Q.-X.J.) and the startup funds from University of Florida. Some of the experiments reported here were performed in a laboratory constructed with support from NIH (grant # C06RR30414 with Dr. Jerry Shay as the PI).

## Methods

### 1. Purification of the recombinant CHGB and its mutants expressed in sf9 cells

We used baculovirus to over-express the protein (Invitrogen). Preparation of other constructs followed the same general procedure as what is described here unless separately stated. The cDNA of the mouse CHGB gene was cloned from a pcDNA3.1 plasmid (a gift from Dr. Barbara Ehrlich at Yale University) into the pFastBac 1 vector (Invitrogen). Cloning was monitored by mapping with restriction endonucleases and PCR-based sequencing. *E. coli* clones with the recombinant bacmids were isolated after the transformation of *E. coli* DH10Bac cells with mCHGB/pFastBac1 plasmid and blue/white-screening of the transformed colonies. The bacmid DNA was used to transfect monolayers of sf9 cells using the CellFECTIN II reagent (Invitrogen). The recombinant viruses were harvested 72-96 hours after transfection (P1), and further amplified twice to obtain higher-titer viruses (P2 and P3). *Sf9* cells in SFM-900 II medium with 2% vol/vol heat-inactivated FBS, and 1x penicillin / streptomycin were infected with recombinant viruses with MOI of one. Cells were harvested 48 hours after infection, and lysed in a buffer made of 50 mM Tris, pH 8.0, 10% glycerol, 5.0 mM DTT, 1.0 mM EDTA, 1.0 mM PMSF, 2.0 % NP40, 1x protease-inhibitor mix (Sigma) and 1.0 μg/ml each of leupeptin, pepstatin and aprotinin. After one hour extraction in the cold room, cell lysates were centrifuged at 100, 000 x g for one hour at 4°C. The clear supernatant was collected and applied directly to pre-equilibrated Q-sepharose FF column (Amersham Biosciences). The column was washed with buffer A (20 mM Tris, pH 8.0, 100 mM NaCl, 2.0 mM beta-ME, 0.050% Triton X-100, 0.5 mM PMSF), and eluted with a three-step gradient of buffer B containing 2.0M NaCl. CHGB-containing fractions were pooled. 25% Ammonium sulfate (final) was added to precipitate CHGB protein. The pellet was collected by centrifugation and dissolved in buffer C containing 20 mM Tris, pH 8.0, 100 mM NaCl, 2.0 mM β-Mercaptoethanol (βME), 0.050 % Triton X-100 of reduced absorbance, and 0.25 mM PMSF. The dissolved mixture was applied to a Ni-IDA column (Amersham Bioscience), washed and eluted with 300 mM imidazole in the buffer. The eluted protein was concentrated and further purified by size-exclusion FPLC using Superose 6 10/30 GL column (Pharmacia Biotech). CHGB fractions were collected and concentrated to ∼5.0 mg/ml using a 30,000 MWCO filter (Millipore).

The CHGB protein concentration was estimated by OD_280_ using a calculated extinction coefficient of 82,405 M^−1^cm^−1^. Purified CHGB was subjected to 10% SDS-PAGE to confirm its purity. For western blot, the protein bands in the SDS-PAGE gel were transferred to a PVDF membrane, immunostained with monoclonal antibodies, and visualized by chemiluminescence (Thermo scientific; SuperSignal^™^ West Pico).

### 2. Expression and purification of individual GHGB fragments from bacteria

Different fragments of mouse CHGB including CHGB-F1, CHGB-F2, CHGB-F3, CHGB-F4, CHGB-Hex, CHGB-Helix2, CHGB-Helix3, CHGB-Cys and CHGB-Cterm (Figure 2A) were individually subcloned into the pGEX-kg (Novagen) vector containing an N-terminal GST-tag. The fusion proteins were over-expressed in *E. coli* BL21 (DE3) host cells (Novagen). After cell lysis and centrifugation, the protein was affinity-purified with Glutathione Sepharose 4B (GE Healthcare). The column was eluted with GSH and the eluted protein was further purified by size-exclusion FPLC in a Superdex 200 column (GE Healthcare) in a buffer containing 20 mM Tris-HCl, pH 8.0, 200 mM NaCl, 1.0 mM EDTA and 1.0 mM DTT with 0.050% Triton X-100.

### 3. Preparation of proteoliposomes

Reconstitution follows closely a published protocol(Lee S. et. al, 2013). 5 or 10 mg of Egg PC lipids (Avanti polar lipids were dried in a glass with Argon gas, vacuum-treated for 1.0 hour, hydrated with autoclaved MilliQ water, and vortexed before being sonicated in an iced water-bath sonicator to make small unilamellar vesicles. N-decyl β-D maltopyranoside (DM; Affymetrix – Anatrace) was added to 40 mM. After 5-6 hours at RT, the detergent/lipid solution became almost completely transparent. Lipid/detergents and proteins in detergents were mixed in a desired protein: lipid molar ratio (PLR), and rocked overnight in a cold room. Next day, BioBeads were added to gradually remove detergents. After reconstitution, the vesicle solution became cloudy, and was aliquoted, flash-frozen in liquid nitrogen and stored at −80°C until experimental use. Control vesicles were prepared similarly without protein. Other proteins were treated in the same procedure even though some of them were soluble and did not get into the vesicles at all.

For vesicle floatation assay, reconstituted vesicles were mixed with 20% Ficoll 400 by 1:3 volume ratio. The mixture (∼0.3 ml) was the bottom layer, above which 10% and 5% Ficoll were loaded sequentially. The gradient was centrifuged at 250,000 x g for 3.0 hours at 4 degrees C. The vesicles migrated to the top 5% layer, and were clearly visible as a narrow band in the gradient. Non-reconstituted proteins stayed at the bottom. The gradients were fractionated for protein detection.

### 4. Recordings of single channel events in bilayer lipid membranes (BLMs)

The CHGB in egg PC vesicles at 0.50 mg/ml in varying PLR of 1:10,000 to 1:2,000 were first tested in 150 to 300-micron bilayer membranes. We observed that immediately after vesicle fusion the membranes were quiet and stable, but in some cases there were sudden changes in a short while (within a few minutes) with multiple high-conductance events, which usually became quiet again after a few tens of seconds. These observations have defied our efforts to record macroscopic currents from many CHGB channels.

Instead vesicles with PLR of 25,000 to 1:10,000 led to small currents (usually < 100 pA at 100 mV) from a handful of channels. The channels remained mostly open in low Vm ([-50, +50 mv]), but frequently closed in higher voltages (Vm >80 or <-80 mV). These channel events often occurred for tens of minutes and then disappeared. We suspected that certain lipid effects might cause such behavior in our recording and will need to use a bSUM to study it (Zheng H. et al, 2016).

PLR 1:10,000 vesicles have on average ∼2 CHGB tetramers per vesicle (assuming 100 nm diameter). Bilayers were prepared as before (Lee S. et al.). Fusion of CHGB vesicles with painted BLMs, was induced by slow water flux from *trans* to *cis* side. After the appearance of channel activities, salt concentration was balanced. To achieve more reliable fusion of vesicles, 0.5 mM CaCl_2_ and 1-2 mM F^−^ ions were introduced in the *cis* side. During the experiments we eliminate those membranes showing negative leak current large than −1.0 pA at 0 mV before salt balancing in the *trans* side because a small currents more positive than −1.0 pA was expected from anion conduction, indicating a tight membrane with little leak. Voltage-clamp mode was used to recording channel activities.

An Axopatch 200B amplifier interfaced with a Windows PC through a DigiData 1322A analogue-to-digital converter was used for recordings. The Axoclamp software (Axon Instrument from Molecular Devices, Inc.) controlled experimental protocols. Clampfit (Axon Instrument) was used to measure currents and events and future data analysis was done in IGOR Pro (WaveMetrics, Inc.). Current recordings were filtered at 1 kHz using a Bessel filter and sampled at 2-5 kHz. The liquid junction potential between the solutions we used was < 0.1 mV. With balanced salt solutions and under 0 mV holding potential, the recording system had a −0.8 to −1.0 pA leak current across a thinned membrane in the absence of any channels. This small negative current was corrected when reversal potential was read out from fitting the I-V curves.

For the recordings made from bilayers under asymmetric Cl^−^ and symmetric K^+^, the *cis* side had 1.5 mM KCl, 150 mM K-isethionate, 10 mM MES-HCl, pH 5.5. The *trans* side started with 15 mM KCl and 10 mM MES, pH5.5, and was balanced with 135 mM K-isethionate after the appearance of channel activities. The liquid junction potential between these two solutions was measured to be close to zero. In these solutions, we recorded no significant outward currents.

We also tested the patching of blebbed membranes fused from reconstituted vesicles without much success. Recordings from giant unilamellar vesicles (GUVs) need a completely different set up, and will be tested in the future for a separate publication.

### 5. Measurement of chloride efflux by a Ag/AgCl electrode

Direct Cl^−^ efflux from vesicles was measured as described previously (Stockbridge et al., 2012). We followed closely what was described in the paper and discussed with Dr. Stockbridge on details. CHGB Liposomes of 1:2,000 PLR (equivalent to ∼10 CHGB tetramers per 100-nm vesicle) were loaded with 300 mM KCl and extruded through a 400-nm membrane filter and passed through a 1.5-ml G-50 Sephadex desalting column in a buffer containing 300 K-isethionate, 25 mM HEPES pH 7.4, 1.0 mM EDTA, 2.0 mM β-mercaptoethanol, 0.2 mM KCl. Vesicles (∼100 microliters; inside KCl and outside K-isethionate) were added to the 1.0 ml recording chamber. The ground Ag/AgCl was connected to the recording chamber through a salt bridge. The recording Ag/AgCl electrode was directly immersed into the recording solution. ∼0.2 mM KCl was added to the recording solution to stabilize baseline. DMSO was added before valinomycin in DMSO stock was introduced at 0.25-1.0 μM to trigger the efflux of Cl^−^. A stirring bar was used to mix well valinomycin with vesicles. Currents were recorded in the whole-cell mode under Vm = 0 mV and at the lowest gain. The data were filtered at 200 Hz and sampled at 2 kHz. Gap-free recordings were made for 45-60 seconds. At the end of the experiments, 50 mM β-octylglucoside (β-OG) was added to release all chloride.

### 6. Measurement of chloride efflux from vesicles by the light scattering assay

For steady-state fluorimetry, we followed closely what was described previously (Stockbridge et al., 2012). Vesicles were loaded with 300-450 mM KCl and 20 mM HEPES at pH 7.4 and were extruded 20 times across a 400-nm membrane filter (Avanti Polar Lipids) before being desalted into a buffer containing 300 mM potassium isethionate. Liposomes were diluted to ∼0.50 mg/ml lipids into 1.0 ml desalting buffer in a stirred cuvette. 1.0 μM valinomycin from 1.0 mM stock solution was added and mixed (∼10-15 seconds) to start the flux. After ∼30 seconds, light scattering was measured at 600 nm inside a Horiba fluoroLog spectrophotometer (HORIBA Scientific Inc.) using a Fluorlog-2 module at UT Southwestern (Jin L, *et al.*, 1999 & Stockbridge et al., 2012). This assay is called the light-scattering-based flux assay. At UF, it was repeated using a Fluoromax-4. Data analysis was performed in Origin.

To titrate the channels per vesicle, PLR varied from 1: 100,000 to 1:1,000. Signals from three independent experiments (different batched of vesicles) were averaged and normalized against the maximal signals. Data were with a Hill-equation:

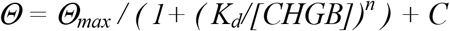

The estimated Hill co-efficient, *n*, for the steady-state measurement is ∼1.4, suggesting that a functional channel needs more than one CHGB subunits. The steady-state measurement suffers from uneven mixing of valinomycin with vesicles due to uneven partitioning at the starting point, and from the slower water diffusion during the relaxation step after a change in vesicle shape.

### 7. Preparation of bacterial EriC transporter

The expression construct was obtained from Dr. Christopher Miller at Brandeis University. The protein expression and purification followed a published procedure (Maduke M et al., 1999). Eric protein was purified using Ni-NTA affinity chromatography and size-exclusion chromatography in a Superdex 200 column (GE healthcare). Protein concentration was measure and reconstituted in egg PC lipids in the same way as described above for the CHGB reconstitution.

### 8. A stopped-flow system to observe the fast kinetics of anion flux

To overcome the shortcomings of the steady-state experiments, we modified the light-scattering based flux assay and used a stopped flow system to achieve quick mixing of equal volumes of valinomycin solution and vesicle solution. An Applied Photophysics SX20 MV stopped flow spectrophotometer (dead time ∼ 2 ms) in Dr. Linda Bloom’s lab was used. It has an observation cell length of 1.6 cm and a mixing chamber of 110 microliters. Right before each experiment, vesicles loaded with 300 mM KCl were exchanged into a buffer containing 300 mM K-isethionate and 10 mM HEPES pH 7.4 and then loaded into one of the injection syringe. The other injection syringe contained 2.0 μM valinomycin in the same buffer. All visible air bubbles were carefully removed. Injection of 55 microliters of solutions from both syringes started the experiments. The reactions were performed at 20°C with a water-bath controlling the temperature of the monitoring cell. Even mixing was achieved within 2 ms. The final mixed solution had 1.0 μM valinomycin, ∼0.4 mg/ml lipids (egg PC) and a varying amount of CHGB protein (PLR). After mixing, light scattering at 610 nm was monitored. Each data point was the average of three consecutive scans, and each experiment was repeated three times using CHGB protein from different batches of purification and reconstitution. Titration of PLRs was performed in a continuous run on the same day to prevent system variation. Because of the slow decay phase after the initial jump, we focused on the first 40 ms in our analysis. To titrate valinomycin, CHGB vesicles of PLR=1: 25,000 was used at varying concentrations of valinomycin. Fitting with a Hill-equation yielded a Hill coefficient of ∼2.6, suggesting strong cooperativity for valinomycin in transporting K^+^ ions to support the fast flow of Cl^−^ ions through CHGB. The IC50 = 650 nM, which is approximately 130 valinomycin in each 100-nm vesicle.

### 9. Negative-stain Electron microscopy of reconstituted CHGB vesicles

Copper grids coated with a thin layer of carbon film were baked at 70°C overnight the day before experiments. After glow discharge, three microliters of reconstituted CHGB vesicles with specific PLRs were loaded. The sample was incubated on a carbon-coated grid at room temperature for 30 seconds before being blotted. After that, the grid was stained with 2.0% phosphotungstic acid (PTA), pH ∼8.0 for 30 seconds. The grid was air-dried before being observed in a JEOL JEM2200FS microscope. Images were taken with a Gatan K2 Direct Electron Detector at 25,000 x with a defocus level of −2.0 microns and a calibrated pixel size of 1.92 Å.

For 3D reconstruction from negative-stain EM images, images were recorded with an electron dose of 20 e^−^/Å^2^ on a 4K × 4K Gatan K2 Summit Direct Electron Detector (Gatan, Pleasanton, CA) in counting mode. 180 images were selected based on the power spectra determined by CTFFIND3 (Mindell JA et al., 2003). 140 images with minimal astigmatism were selected for particle picking. The particles were selected using the Boxer module in EMAN 2 (Tang G, *et al.* (2007). A total of ∼5,400 particle images were manually selected. An initial model was generated by angular reconstitution in IMAGIC 5 (van Heel M et al., 1996) and finally refined in SPIDER (Frank J, *et al.*, 1996 & Jiang QX et al., 2003). The final map was calculated from ∼3,000 particle images at a nominal resolution of 30Å. The handedness of the map was tested from images from +15 and −15 degrees as we did for the C3PO negative stain map (Llaguno MC, *et al.*, 2014). The small, compact size of the dimer made us less confident in the handedness at this point. A high-resolution map will be used to further examine chirality.

### 10. CryoEM study of CHGB dimers

Quantifoil R2/2 grids (Quantifoil Micro Tools GmbH, Jena, Germany) were coated with a thin carbon film (∼2–3 nm). The ChemiC (Ni-NTA) grids were prepared as described before (Anonymous, 1986). 3.0 μl of purified CHGB in detergents was loaded. After incubation for ∼15 minutes in a wet-chamber of >90% humidity, the grid was blotted inside a Vitrobot and plunge-frozen into liquid ethane bathed in liquid nitrogen (FEI, Hillsboro, OR). After screening in a JOEL2200, good specimens were imaged at HHMI Janelia Farm Research Campus. A Titan Krios microscope equipped with a 4K × 4K Falcon 2 Director (no movie function at the time) was used. The scope was operated at 300 kV and was equipped with a *Cs* corrector. Automatic data collection was run by a proprietary software package, EPU (FEI, Hillsboro, OR). Images were taken under a defocus of −2.5 to −4.0 microns at a magnification of 37,000 ×, which gave rise to a calibrated pixel size of 1.89 Å at the specimen level.

Because the CHGB dimers were quite small, we scanned many areas for good recognition of the particles. Only a small dataset was successfully built, which came from ∼300 of 4K x 4K images. These images all displayed good Thon rings to a resolution of ∼ 6.0 Å with minimal astigmatism and defocus values ranging from −1.0 to −4.0 μm and showed visible particles. 24,086 particles were picked manually and extracted in 196×196 Å^2^ boxes. The low-resolution negative-stain map was used as the reference for 3D refinement. Five rounds of 2D classification into 50 distinct classes were done with the program RELION 1.3 (Scheres SH., 2012). The classes with well-defined particles were selected. 3D classification was performed with these selected particles into five classes. Two classes showing higher resolution features were selected for further refinement. The 12,123 particles that were assigned to these two classes were subjected to 5 additional rounds of refinement using a high-resolution frequency limit of 6 Å. A soft mask was introduced to redo the 3D classification and remove ∼45% of the particles. The final map was calculated from ∼6,900 particles and the estimated resolution was 9.8 Å by using a threshold of 0.143 to the gold-standard Fourier Shell Correlation (van Heel M et al., 2005 & Rosenthal PB et al., 2003). The map was sharpened by applying a negative B-factor of −75 Å^2^.

### 11. Measuring Ca^2+^ efflux from reconstituted vesicles

Proteins (CHGB, its mutants or IP_3_R) were reconstituted into vesicles in a PLR of ∼1:5000 in a buffer made of 20 mM HEPES, pH7.5, 100 mM NaCl, 1.0 mM EDTA and 2.0 mM β-ME (high pH) or a buffer made of 20 mM MES, pH5.5 (low pH). To load CaCl_2_, 1.0 mM CaCl_2_ was added to the buffer. Right before each experiment, freshly prepared lipid vesicles 10 mg/ml) was extruded by 20 stokes through a membrane filter with an average pore size of 100 nm (Avanti polar lipids). A PD-10 desalting column (Sigma) was used to change the vesicles into a buffer containing 100 mM NaCl and 20 mM HEPES, pH 7.4, 1.0 mM EDTA and 2.0 mM β-ME. Chelex 100 was used to treat the buffers in order to remove residual calcium ions. Vesicles out of the column (∼3.0 mg/ml lipids) were diluted to 0.2 mg/ml into the external buffer inside a quartz cuvette with constant stirring. A Ca^2+^-sensitive fluorophore, Indo-1, was added to 1.0 μM. The efflux, if any, was initiated by adding 0.5 μM valinomycin. Indo-1 fluorescence at 410 nm was measured in a Horiba fluoroLog spectrophotometer (HORIBA Scientific Inc.) using the Fluorolog-2 module and an excitation wavelength of 330 nm.

For the IP_3_R-containing vesicles, the protein was purified from rat cerebellum as reported before (Jiang QX et al., 2002), and reconstituted in egg PC in the presence of 2.0 mM CaCl_2_. The efflux of calcium was initiated by adding 1.0 μM IP_3_ at different time points. At the end of the experiments, 30 mM β-octylglucoside was added to disrupt the liposomes and release all calcium ions to determine the maximum signal.

### 12. Knockdown of CHGB in neuroendocrine cells and rescue by overexpressing CHGB or CHGBΔMIF

INS-1 (832/13) cells were seeded one day before transfection at 40% confluency. Cells were transfected with RNAiMAX (Life Technologies) that was mixed with varying amounts of siGENOME non-targeting (D-001210-01-05; CTL siRNAs or scRNAs) and CHGB siRNA-SMART pool (M-099320-01-0005; CHGB siRNAs; or the 2^nd^ and/or 4^th^ siRNA) from Dharmacon.

To examine the knockdown effect, we lysed the cells for western-blot. 96 hours after transfection, cells were collected using 10 mM EDTA in 1 x PBS. Extracellular EDTA was removed from the cells by washing them twice with PBS. The cells were changed into a lysis buffer containing 50 mM Tris, pH8.0, 10 mM NaCl, 1.0 mM EDTA and 1.0 mM DTT, and went through three freeze-thaw cycles between liquid N_2_ temperature and 37 degrees C (Zheng K. et al., 2014). Afterwards, cell lysates were incubated in ice for 30 min and centrifuged at 18,000-x g for 30 min to remove cell debris. Concentration of the released protein was measured using Nano-Drop 2000c spectrophotometer (Thermo Scientific) or more accurately using the Bradford method (Bio-Rad). Roughly 20 μg of protein from cell lysates was used for each lane in a SDS-PAGE gel for western blotting with anti-CHGB antibody (C-19: catalog #: sc-1489: Santa Cruz Biotechnology, Santa Cruz, CA, USA). (HRP)-conjugated donkey-anti-goat 2^nd^ antibody (Santa Cruz) was used for detection using the Super Signal West Pico chemiluminescent substrate (Thermo Scientific).

### 13. Detection of insulin and proinsulin in INS-1 cells

INS-1 cells were transfected with 100 nM control siRNAs or CHGB siRNAs in a 6-well plate and fed with fresh medium after 48 hours. 96 hours after transfection, cells were collected using EDTA and lysed by freeze-thaw cycles as described above. An anti-insulin antibody (L6B10 from Cell Signaling) was used to detect insulin and proinsulin. The band intensity in western blot images was measured in ImageJ (Schneider CA. et al., 2012).

For the rescue experiments, cells transfected with CHGB siRNAs were incubated for 48 hours, and then transfected with a pcDNA3.1 plasmid carrying the genes for the wild-type CHGB or its deletion mutant. After two more days the cells were collected for western blotting analysis. A 12% tricine-SDS-PAGE gel was used for separating insulin and proinsulin. The internal loading control was also used to correct for small variations.

### 14. Western blot detection of V-ATPase subunit A2

Cells were treated the same as above. The collected cells were lysed by freeze-thaw cycles. Roughly 20 mg of lysate protein were separated by SDS-PAGE and detected by western blot using an anti-ATP6V0A2 antibody (Abcam).

### 15. ELISA assay for detecting dopamine released from PC-12 cells

On day 0, 3 x10^5^ PC-12 cells were seeded into each well of a 6-well culture plate. CHGB siRNAs transfection was performed in Day 1. On day 3, cells were transfected with the plasmids containing cDNAs for wild-type CHGB or CHGBΔMIF. On day 5, cells were prepared for analysis. The cells without rescue were fed with fresh medium on day 3. The dopamine secretion test was conducted as previously described with slight modifications (Shoji-Kasai Y. et al., 2002). The cells were washed twice in a low (basal) K^+^ buffer (20 mM HEPES-NaOH, pH 7.4, 140 mM NaCl, 2.5 mM CaCl_2_, 1.2 mM MgCl_2_, 5.0 mM glucose, and 4.8 mM KCl). Equal number of cells were then treated in parallel with a low K^+^ buffer (4.8 mM KCl) as control or a high K^+^ buffer (40 mM KCl) for 11 min at 37 °C. Afterwards, the media were collected and immediately centrifuged at 10,000 × g for 20 s at 4 °C. The dopamine concentrations in the supernatants were measured using a commercial ELISA kit (Dopamine ELISA Assay kit from Eagle Biosciences). We performed these experiments four times and did not observe obvious toxicity to the PC-12 cells due to the siRNA or cDNA transfection. The measured dopamine was thus normalized against that from the cells transfected with CTL siRNAs. The amount of total protein contained in the cell lysates was measured to confirm that the measured dopamine was released from approximately the same number of cells treated in four different conditions.

### 16. pH measurements in intracellular acidic compartments

First we tested whether the lysosensor DND-160 would be suitable for our measurement. INS-1 cells were transfected to express Syncollin-pHluorin fusion protein (Fernandez Na. et al., 2011) (a construct provided by Dr. Herbert Y. Gaisano in the Department of Medicine at University of Toronto, Canada). 48 hours after transfection, cells were washed with the pre-warmed physiological medium and incubated with 0.5 μM DND-160 for five minutes at 37°C. Afterwards, cells were rinsed quickly and imaged in a Zeiss LSM 780 (63 x oil-immersion lens) confocal fluorescence microscope using 485 nm excitation and 510 ±10 nm emission for pHluorin and 405/510 nm for DND-160. At UF, a Zeiss LSM 800 confocal microscope was used instead.

INS-1 or PC-12 cells were prepared as described above. The imaging was always done 96 hours after siRNA transfection. Immediately before imaging, cells were washed with a physiological medium (10 mM HEPES, 120 mM NaCl, 4.8 mM KCl, 2.5 mM CaCl_2_, 1.2 mM MgCl_2_, 24 mM NaHCO_3_, 3.0 mM Glucose, pH 7.4) and then incubated with 1.0 μM DND-160 for 5 min at 37 degrees C. 0.50 μM Bafilomycin was added. Excess dye was washed away with the physiological medium and images of the cells were captured in a microscope. For double excitation experiment, a 40-x Quartz objective was used and the dye was excited at 340 nm or 380 nm and the emission was recorded at 510 nm. We used a home-assembled Fura-2 imaging system in Dr. Jen Liou’s laboratory at UT Southwestern Medical Center. For every granule from each image pair, a circular mask of the granule size was used to read the average density. Right in proximity the same mask was used to read out background signal from four blank areas. The averaged background signal was subtracted, generating the final intensity for the selected granule. The paired images were analyzed side-by-side so that the same areas were selected. A 340 nm / 380 nm ratio was calculated (dual excitation). For dual emission, the cells were prepared the same way and imaged under a LSM 780 (or LSM800) upright confocal microscope (Apochromat 63x/1.40 Oil DIC objectives). Images were collected using 405 nm excitation and emission at 510 nm (yellow) and 485 nm (Blue) with 20 nm apertures. The ratio of 510/485 for individual granules was calculated. The standard curves for dual excitation and dual emission were prepared in the same imaging systems using the DND160 in buffers at different pH values. The standards curves were used to avoid cell-to-cell variations. Because of the reported difficulty in controlling pH inside the granules for calibrating pH *in situ* (Deriy LV, et al., 2009), we only used the *in vitro* calibration. The histogram distribution of the measure pH values therefore reflected the distributions of the ratiometric measurements based on the calibration and may not reflect very accurately the *in situ* pH, and positive and negative control experiments were used to demonstrate the suitable application of the measurements as a reliable quantification of the alterations in intragranular pH. A large number of granules were randomly selected from multiple cells in each condition in order to overcome small variations from one cell to another and from granules of different sizes or in different states and derive statistically significant and objective comparison.

### 17 Proteomic analysis of proteins

Trypsinization and mass spec analysis of the protein bands were performed by the Protein and Peptide core facility at UT Southwestern Medical Center. The N-terminal sequencing of the purified fragments eluted from the SDS-PAGE was performed by the facility. Standard procedures at the core facility were used. After my lab was relocated, the Proteomics core at UF ICBR performed the same services.

The excised bands from the Coomassie-blue-stained SDS-PAGE gels were submitted for identification. The gel bands were de-stained with 1.0 ml of 50 mM ammonium bicarbonate, pH 8.0/acetonitrile (1:1, v/v). Each sample was reduced with 40 mM DTT, alkylated with 100 mM of iodoacetamide, and trypsin-digested. Trypsin-digested peptides were desalted with C18-Ziptip (Millipore, Billerica, MA, USA) A hybrid quadrupole Orbitrap (Q Exactive Plus) MS system (Thermo Fisher Scientific, Bremen, Germany) interfaced with an automated Easy-nLC 1200 system (Thermo Fisher Scientific, Bremen, Germany) was used. Samples were loaded into an Acclaim Pepmap 100 pre-column (20 mm × 75 μm; 3 μm-C18) and separated in a PepMap RSLC analytical column (250 mm × 75 μm; 2 μm-C18) with a flow rate of 250 nl/min using a linear gradient from solvent A (0.1% formic acid (v/v)) to 25% solvent B (0.1% formic acid (v/v), 19.9% water (v/v) and 80 % acetonitrile (v/v)) for 80 min, and then to 100% solvent B for additional 15 min.

The spectrum library was produced in a data-dependent mode with survey scans acquired at a resolution of 70,000 at 200 m/z. The mass spectrometer was operated in MS/MS mode scanning from 350 to 1800 m/z. Up to 10 most abundant isotope patterns with charges of 2-5 from the survey scan were selected with an isolation window of 1.2 Th and fragmented by high energy collision dissociation (HCD) with normalized collision energies of 28. The maximum ion injection times for the survey scan and the MS/MS scans were 250 ms, and the ion target values were set to 3.0E6 and 1.0E6, respectively.

A software package, Scaffold (v4.7.5, Proteome Software Inc., Portland, OR), was used to validate MS/MS-based peptide and protein identifications. Identified peptides were accepted if they were established at greater than 93.0% probability by the Peptide Prophet algorithm (Keller A, et al., 2002) with Scaffold delta-mass correction. Protein identifications were accepted if they could be established at greater than 97.0% probability and contained at least 1 identified peptide. Protein probabilities were assigned by the Protein Prophet algorithm (Nesvizhskii Al, et al., 2003). Proteins that contained similar peptides and could not be differentiated based on MS/MS analysis alone were grouped to satisfy the principles of parsimony. Proteins sharing the identified peptides were grouped into clusters.

### 18. Estimated ion flow rate through the CHGB conductance requires a fast-conducting channel, not a slow-acting transporter

The main difference between a transporter and a channel is the flux rate. Even for the fastest known ion transporter (Cl^−^/HCO_3_^−^ transporter), its turnover rate (up to 10^5^ per second) would still be several orders of magnitude slower than that (∼10^7^ per second) of a channel. The stopped-flow-based flux assays can provide a good estimate of the flux rate that is limited by the maximum flow through valinomycin molecules. Titration of valinomycin concentration in Fig 5H suggested that ∼200 valinomycin molecules per vesicle were needed to generate a significant signal within the 2-ms fast mixing. We assume that almost all valinomycin molecules partition to lipid membranes due to their hydrophobicity. Valinomycin shows a turnover rate of ∼10^4^ per second at room temperature in the absence of a transmembrane electrostatic potential. Thus, the estimated K^+^ flux per 100 nm vesicle is ∼ 2 × 10^6^ per second, or ∼4 × 10^3^ in 2 ms, which alone is not enough to release a significant fraction of ∼10^5^ K^+^ ions from each 100-nm vesicle without a transmembrane potential.

In the very beginning of mixing two solutions the mixture had a lipid concentration of ∼0.4 mg/ml, a PLR varying from 1: 100,000 to 1: 1,000 (molar ratio), and a [valinomycin] of 1.0 micromolar. With 300 mM KCl inside the vesicle and K-isethionate outside, the Nernst potential was infinite, but with a small leak of Cl^−^ to the outside, an electrostatic potential was established. The Nernst equation was related to the charging of the vesicles after ∼580 chloride ions moved outside. A numerical solution was obtained using MATLAB. The following shows the parameters for individual vesicles.

|  |  |
| --- | --- |
| Average diameter: | 100 nm |
| Capacitance coefficient: | $\sim 1 \mu F/cm^2$ |
| Surface area of each lipid (PC) molecule(White SH. et al., 1985): | $\sim 70 \text{ \AA}^2$ |
| Average number of lipid molecules per 100 nm vesicle: | $\sim 8 \times 10^4$ |
| Average vesicle concentration (0.4 mg/ml lipid): | $\sim 5.0 \text{ nM}$ |
| Volume inside a 100 nm vesicle: | $\sim 0.41 \text{ aL}$ |
| Number of $K^+$ or $Cl^-$ ions per 100 nm vesicle: | $7.4 \times 10^4$ |
| Number of $Cl^-$ ions moved to reach a stable Nernst potential: | $\sim 580$ |
| Average $[Cl^-]$ outside when a stable Nernst potential was reached: | $\sim 2.6 \mu M$ |
| Steady-state Nernst potential before valinomycin addition: | $\sim +295 \text{ mV}$ |

Please note that the Nernst potential became stable once shifted away from the initial infinity. From these estimates, a strong Nernst potential inside would drive the valinomycin transport of K^+^ ions out. An electrostatic driving force for K^+^ efflux at the first 2 ms can be expressed as a Correction Factor -- *exp(zδFV/RT), which is* ∼8.9 × 10^4^ at +295 mV, where *δ* is a factor for effective conversion of the electrostatic energy into the mechanic motion and is assumed to be unity here under the consideration of no significant energy loss due to either partial charge loss or charge delocalization during the movement of the K^+^/valinomycin across the bilayer. Under this consideration, the initial K^+^ flux rate through ∼200 valinomycin molecule in each vesicle could be as high as ∼1.8×10^8^ per ms at +295 mV, which is the peak rate. Once the K^+^ and Cl^−^ions started to flow out, correction factors quickly decays when the vesicular potential drops to below 180 mV. Such a high initial rate would be enough to quickly dump most of the ∼10^5^ K^+^ ions inside each vesicle in < 1 ms, and cause a significant, sudden decrease in osmolality and thus a sudden collapse of the vesicles before water diffusion could follow the change. The water movement was found to be in the tens of ms range. This explanation would also account for the sensitivity of the light-scattering signal to [valinomycin]. Moreover, for the fast removal of KCl from the interior of each vesicle, the efflux of chloride ions had to match that of the potassium ions. A lower limit is ∼10^7-8^ per second. Possible movement of water molecules accompanying the movement of Cl^−^ only accounts for a tiny fraction of water molecules inside each vesicle.

Our calculation made ta testable prediction that a slight increase in [Cl^−^] outside of the vesicles would diminish the flux signal by influencing the Correction Factor. Our data showed that 2-5 mM KCl in the extravesicular side almost completely abolished the light-scattering signal. It means that when the initial transmembrane Nernst potential drops to < +150 mV inside (0.875 mM KCl outside), the correction factor of *exp (zδFV/RT)* decreases by ∼290-folds such that the average KCl efflux rate would become significantly slower than what’s needed to generate a sudden drop in osmolality. Our experimental data confirmed our prediction.

Based on the titration of PLRs in CHGB vesicles in the stopped-flow assay, the average number of CHGB subunits per vesicle needs to be above 4 per vesicle for generating a fast signal, which is likely equivalent to one channel, and 6-8 CHGB subunits per vesicle are sufficient to reach a maximum signal because of the random Poisson distribution (See the next section). That is, on average 1-2 CHGB tetramers are sufficient to achieve the fastest efflux estimated. A flux rate of ∼10^4-5^ ions per ms per molecule can only be achieved by fast diffusion through an ion channel with a significant single channel conductance. It can not be done by even the fastest known transporter, a Cl^−^/HCO_3_^−^ transporter, with a turnover rate of ∼ 10^2^ per millisecond. From the estimated single channel conductance of CHGB under difference ion conditions, a conductance level of 58 pS (in the presence of low [Cl^−^]) is capable of conducting ∼1.1 × 10^5^ ions per millisecond, quite suitable for generating a sudden drop of osmolality.

The fast mixing and even partitioning of valinomycin with the vesicles in the stopped-flow mixing cell and the relatively lipophilic nature of valinomycin makes it possible for us to assume that it takes no free energy for valinomycin to diffuse through the aqueous phase in an estimate velocity of ∼1-2 μm/ms via random walk, and become enriched in bilayer membranes. The efflux of a small amount of K^+^ ions carried by valinomycin is followed by the efflux of Cl^−^ ions through the CHGB channels such that the positive potential inside the vesicles would fall quickly from the starting +295 mV, following the exponential curve. The change in potential will slow down the efflux of K^+^/Cl^−^ and eventually stop further change in osmolality and vesicle shape.

### 19. A Poisson distribution of CHGB subunits among vesicles and the estimated stoichiometry of functional channels

For random reconstitution, a Poisson distribution of the CHGB dimers among individual vesicles is expected. For example, a PLR=1:10,000 yields an average number (λ) of CHGB monomers per 100-nm vesicle to be ∼8. This distribution allows us to evaluate the fraction of vesicles that might have enough CHGB subunits (N) to form a functional channel, assuming that functional channels are tetramers, or higher-order oligomers (N = 4, or more).

Given that 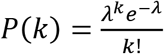 represents the probability of vesicles containing on average *k* copies CHGB monomers, where *k* is a discrete number (0, 1, 2, 3, …) and λ the average number of CHGB monomers per vesicle that is set by adjusting the PLR of the CHGB vesicles, we calculated the fractions of vesicles that would each have less than N copies of CHGB monomers with N = 2, 3,…, 8. The theoretical predictions are here compared with experimental data in the last column (green).

| | | Fraction of vesicles have fewer than N monomers, $P(k < N)$ | | | | | | | Stopped-flow data |
| --- | --- | --- | --- | --- | --- | --- | --- | --- | --- |
|  |  | 2 | 3 | 4 | 5 | 6 | 7 | 8 |  |
| $\lambda$,<br><CHGB><br>/ vesicle | 0.8 | 0.81 | 0.95 | 0.99 | 1 | 1 | 1 | 1 | 1 |
|  | 1.67 | 0.503 | 0.765 | 0.911 | 0.972 | 0.993 | 0.998 | 1 | 0.912 |
|  | 2 | 0.406 | 0.677 | 0.857 | 0.947 | 0.958 | 0.983 | 0.996 | 0.846 |
|  | 4 | 0.092 | 0.238 | 0.434 | 0.629 | 0.785 | 0.889 | 0.949 | 0.481 |
|  | 8 | 0.003 | 0.014 | 0.042 | 0.1 | 0.191 | 0.313 | 0.453 | 0.19 |

We consider the following assumptions. 1) When N monomers are available in a vesicle and sufficient to join together to form a channel, the assembly is nearly 100% efficient. Based on the biochemical data, the CHGB protein was fairly stable and our reconstitution procedure was able to incorporate all protein into vesicles in a preferred orientation; 2) One channel suffices to conduct enough Cl^−^ from the interior of a vesicle within 1-2 ms. The conductance of CHGB with 300 mM Cl^−^ would be even larger than 125 pS; 3) With assumption 1, all vesicles with less than N copies of CHGB monomers will have no channel and will not contribute to the light-scattering change. These together will form a non-functional fraction with a probability of *P(k<N)*; 4) Those vesicles with more than N monomers will form at least one functional channel, and the surplus monomers (very likely dimers as the basic units) will not interfere with the function of the assembled channels. Because we are using a large number of vesicles for our experiments, the statistical average will likely overcome the experimental variations in vesicle size, completeness in the CHGB insertion from the extravesicular side into individual vesicles, freedom of CHGB dimers to diffuse and interact with each other in vesicles, relative ratio of individual lipid types in the egg PC mixture, diffusion time for valinomycin molecules to arrive at membranes, and degree of reaching complete mixing of two equal volumes within the 2 ms dead time. We think that it is relatively reasonable that these assumptions will be satisfied.

Our experimental data in the stopped-flow measure are listed to the last column. When a least squares analysis was used to compare the differences between the experimental data and the theoretical predictions, the best fit is N=4, which is better than N=5, but much better than N=2, 3, 6, 7 or 8. The agreement of the predictions from Poisson statistics and the experimental data appears not co-incidental, especially at the lower range of average CHGB monomer per vesicle (λ <= 4), where we expect that the assumption of random distribution is better satisfied. Considering the dimers being the dominant species in detergents and the tetrameric form observed in the chemically cross-linked fractions, our data support the tetramers as functional channels, not the pentamers or other oligomers.

### 20. Isolation of pancreatic islets from wild-type and CHGB knockout mice

Following a UF IACUC protocol (#201709886), we imported the CHGB knockout mice from the InfraFrontier GmbH in Germany. The heterozygous CHGB C57BL/6 stain was #EM: 10088 in the EMMA deposit with a common strain name of CEPD0073_3_D01 and a detail strain description of **C57BL/6NTac-Chgb^tm1(EGFP/cr/ERT2)Wtsi^/WtsiIeg**. The URL for the deposit site is: https://www.infrafrontier.eu/search?keyword=CHGB. The genome-wide mutants were first produced by Wellcome Trust Sanger Center and EMMA is distributing them. The mice were first raised from frozen sperms and shipped to us as five heterozygous (3 males and 2 females). After quarantine at the UF Animal Care Service, the mice were raised in the Animal Breeding facility. Genotyping of all mice were performed by PCR amplification of DNAs extracted from tail nips. Information for the primers used for PCR was obtained from EMMA. West blotting to detect full-length CHGB was performed from processed tissues including liver, pancreas, spleen and kidney, etc. We kept three pairs of heterozygous mice for continuous breeding. Every time, we used two wild-type (one male and one female) and two homozygous (one male and one female) for surgical preparation of pancreatic islets. The mice were dispatched by the Animal Care Service on the day of experiments and sacrificed on the same day.

Surgical removal of pancreas from C57BL/6 mice was followed using a standard protocol (Pinar Yesil, et al., 2009 & Ablamunits, V. et al., 1999). After euthanizing with CO_2_, the mouse was laid in supine position and sanitized by spraying with 70% ethanol in H_2_O. The mouse abdominal cavity was surgically opened to expose small intestine, common bile tubes, pancreas, etc. Cardiac aorta was cut to ensure complete animal euthanization. Under a dissecting microscope, the small intestine of the mouse was repositioned so that the common bile duct formed a perpendicular line with the head of the mouse. The common bile duct was clamped in a position as close as possible to the small intestine. 2.0 ml Liberase TL (Roche) solution at 0.9 unit/ml concentration was injected slowly using a 27G1/2 Gauge needle via a nylon tubing through bile duct. After that, the pancreas was gently removed from the animal and stored in a 50 ml conical tube in ice. Usually all four mice were performed one at a time. Their genetic phenotype and gender was marked on the conical tubes. After the surgery the mice carcasses were properly disposed.

After surgery, the conical tubes containing the pancreas were incubated in a water bath at 37°C for 14 minutes. 25 ml HBSS with Calcium and Magnesium was added into each tube, and pancreatic tissues were disintegrated by pipetting the solution up and down. The tubes were transferred immediately to a centrifuge at room temperature for a quick spin at 200x g for 2 minutes. The pellets were washed with HBSS four times. The pellets were gently resuspended in 10.0 ml DMEM medium and the tissue suspension was poured into a 15mm x 100 mm polystyrene petri dish, where the islets were hand-picked and transferred into fresh medium. The islets were incubated at 37°C with 5% CO_2_ for 48 hours before used for experiments. The intragranular pH measurements in beta-cells of isolated islets followed the same procedures we used for cultured cells. After washing with HBSS, the cells were stained with DND-160 and washed for imaging under a Zeiss LSM-800 confocal microscope. Pairs of images were obtained in tandem together with the DIC image. To avoid the problem from uneven DND-160 staining, the focal level was limited to the first one or two layers of beta-cells under the islet surface. Data analysis followed what was described for INS-1 and PC-12 cells.

**Supplementary Table S1.**
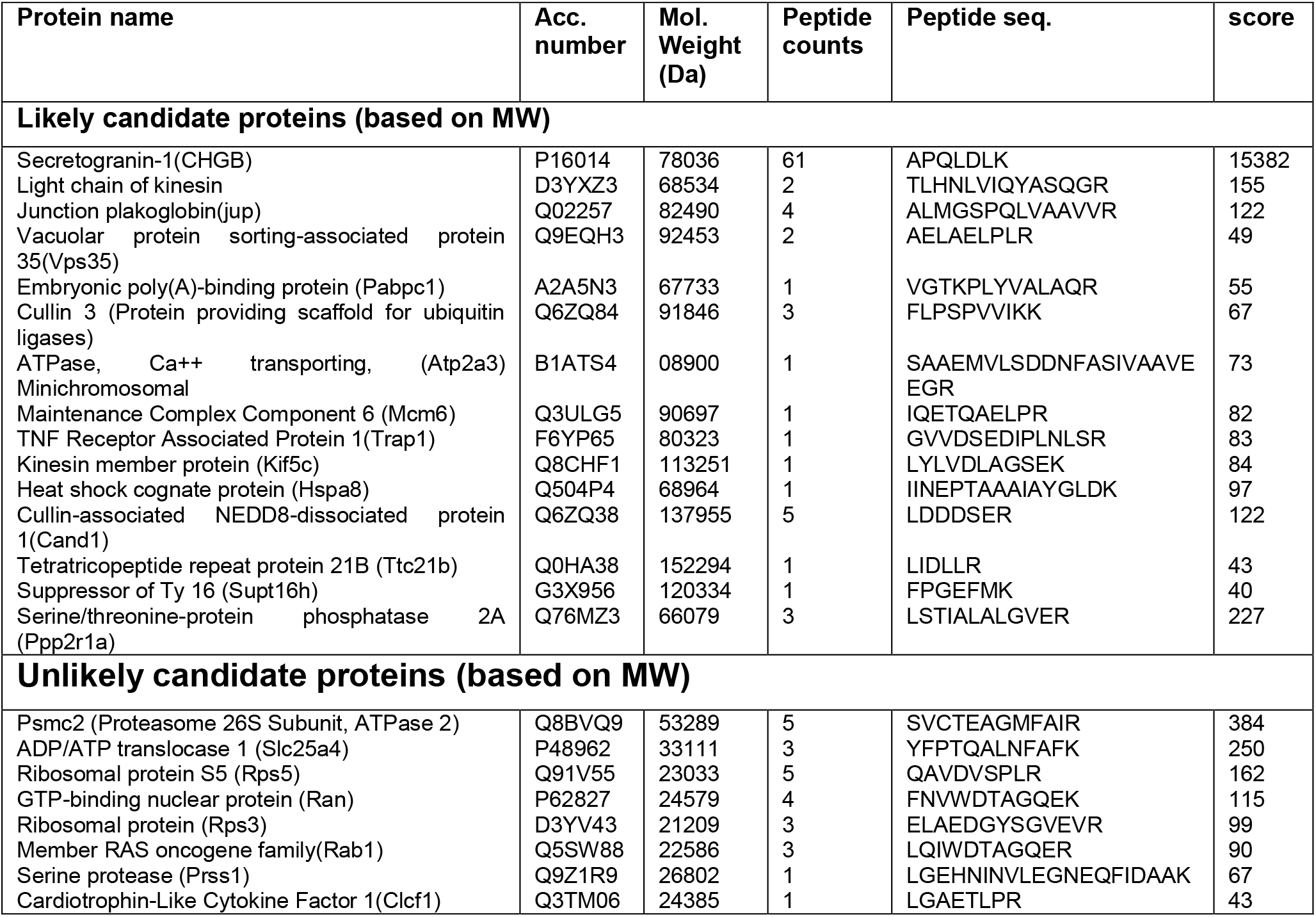
Proteins identified by Mass Spec and proteomic analysis of the full-length CHGB band.

**Supplementary Table S2.**
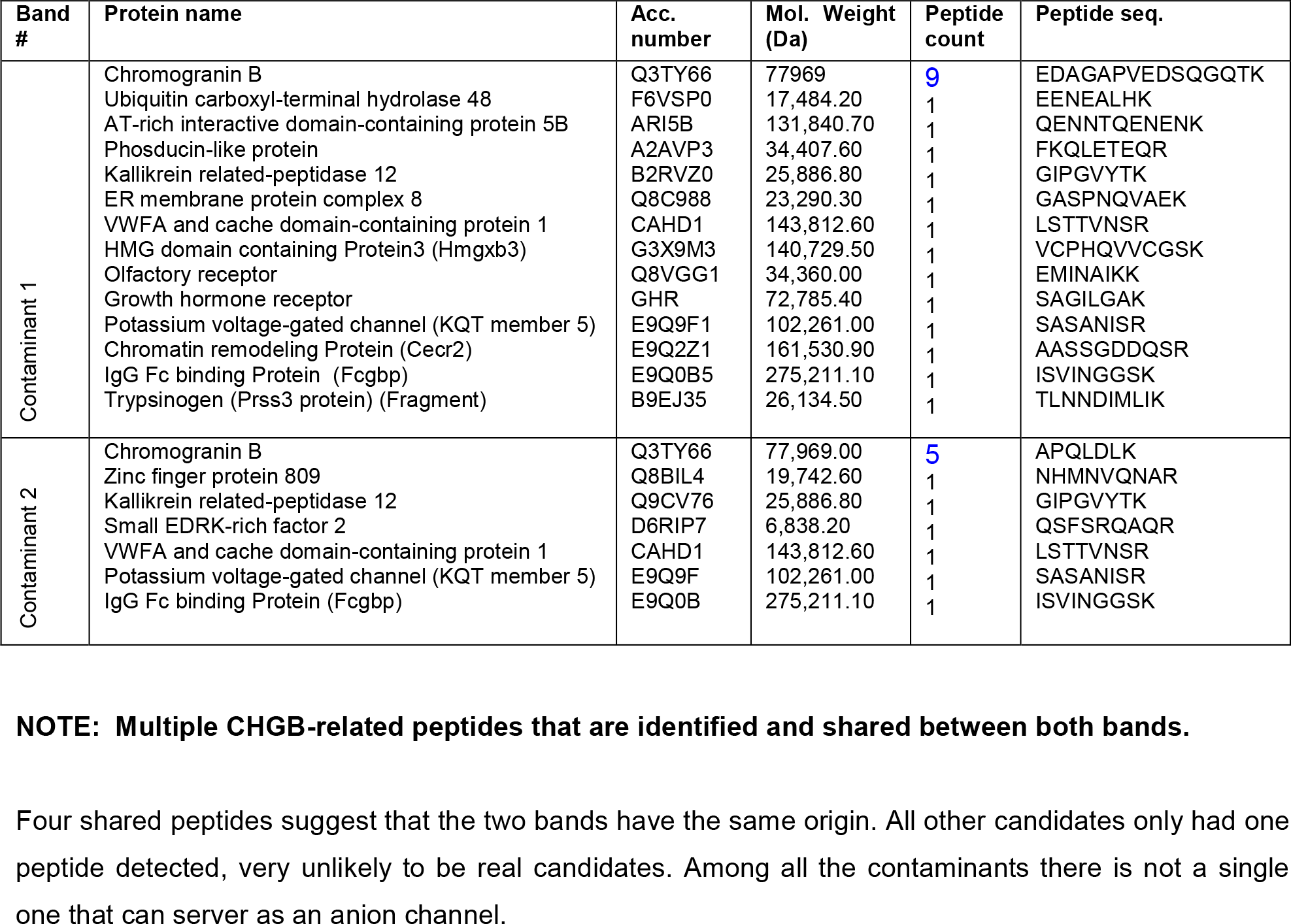
Proteins identified in two contaminating bands

**Supplementary Figure S1.**
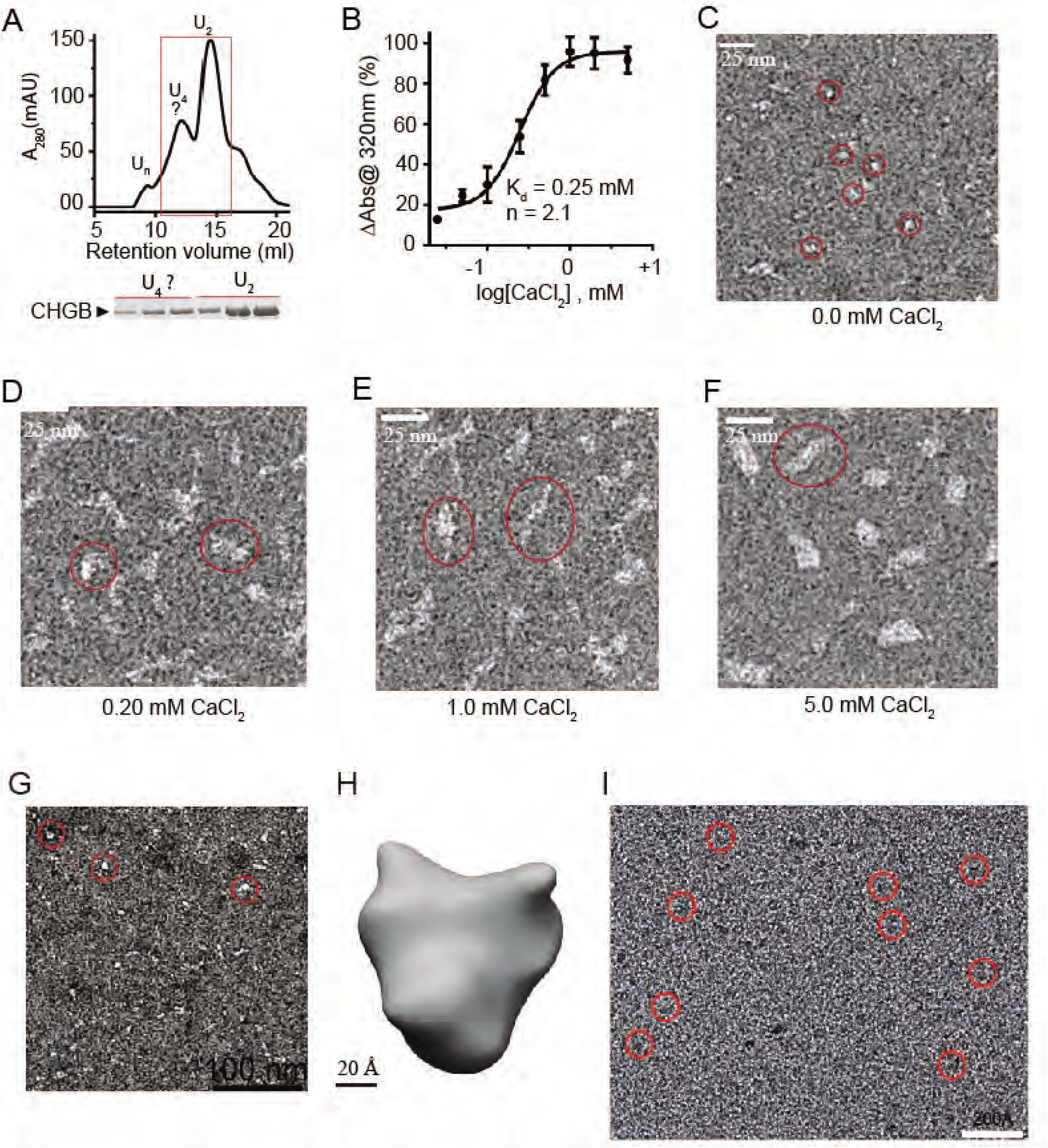
Calcium induced CHGB aggregation and single particle analysis of CHGB dimers. **A)**. SEC profile of partially purified CHGB in a Superose 6 column. The dimer peak (U_2_) was collected for further preparation. (**B**) 20 micrograms CHGB incubated with different [CaCl_2_] for 30 min at RT and OD was measured at 320 nm. Change in absorbance (*ΔAbs* @320nm) was plotted against log [CaCl_2_] and fitted with a Hill equation:

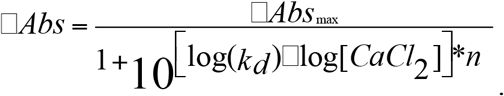 Fitting (black line) yielded *k_d_* ∼ 0.25 mM, a Hill coefficient *n* ∼2.0, suggesting the apparent positive cooperativity in the Ca^2+^-stimulated aggregation. Error bars represent *s.d.* (n=3). (**C-F**) EM images of negatively stained (2% PTA) CHGB incubated with different [CaCl_2_]. Red circles highlight some of the CHGB dimers (C) or aggregates (D-F). (**G**) Images of negatively-stained CHGB dimers in detergents. The protein concentration was 0.1 mg/ml. The stain was 2.0% PTA/KOH pH 8.0. (**H**) A 30 Å negative-stain map calculated from 3,200 particles. Scale bar: 2.0 nm. (**I**) Typical cryoEM images of the CHGB dimers on the surface of a ChemiC-Ni-NTA grid. Some of the CHGB particles are highlighted by red circles. The protein is in black from the raw images.

**Supplementary Figure S2.**
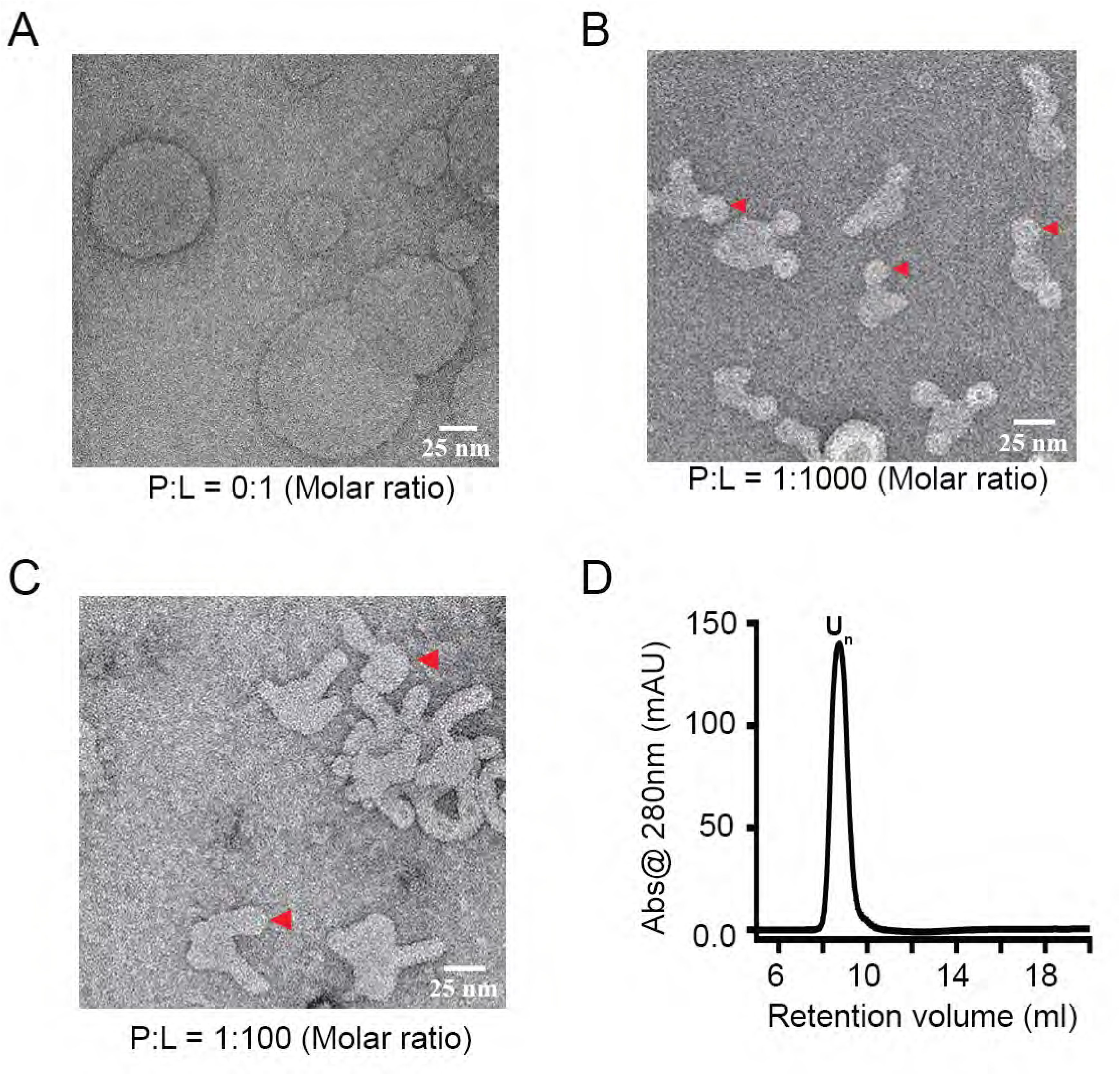
Membrane insertion of CHGB introduces positive curvature in the bilayers. Purified CHGB protein was reconstituted in lipid vesicles made of egg PC. Empty vesicles were prepared in the same way without CHGB. The vesicles were diluted by 5 fold in a buffer containing 20 mM Tris-HCl, pH 7.4, 100 mM NaCl, 1.0 mM EDTA, and 2.0 mM beta-ME before being loaded on carbon-coated grids and stained with 2.0 % PTA (phosphotungstic acid / KOH, pH 8.0). Typical images of empty vesicles (**A**), CHGB reconstituted vesicles at 1:10 protein/lipid weight ratio (1 : 1,000 in molar PLR or ∼40 CHGB dimers per 100 nm vesicle) (**B**) and CHGB reconstituted vesicles at 1:1 protein/lipid weight ratio (1:100 molar PLR) (**C**) are shown. The experiments were repeated more than three times with very similar results. Budding compartments with positive curvature are marked with red arrowheads in **B and C**. In B, the nanospheres are marked at the tops of nanotubules. (**D**) SEC profile of high-order CHGB oligomers (larger than tetramers) from detergent extracts of the CHGB vesicles. The molecular weight of the peak fraction, ∼1.0 MDa, was estimated by using a broad-range MW standard. The extracted proteins were re-run in an SEC column in the presence of 0.1 mg/ml egg PC in detergents.

**Supplementary Figure S3.**
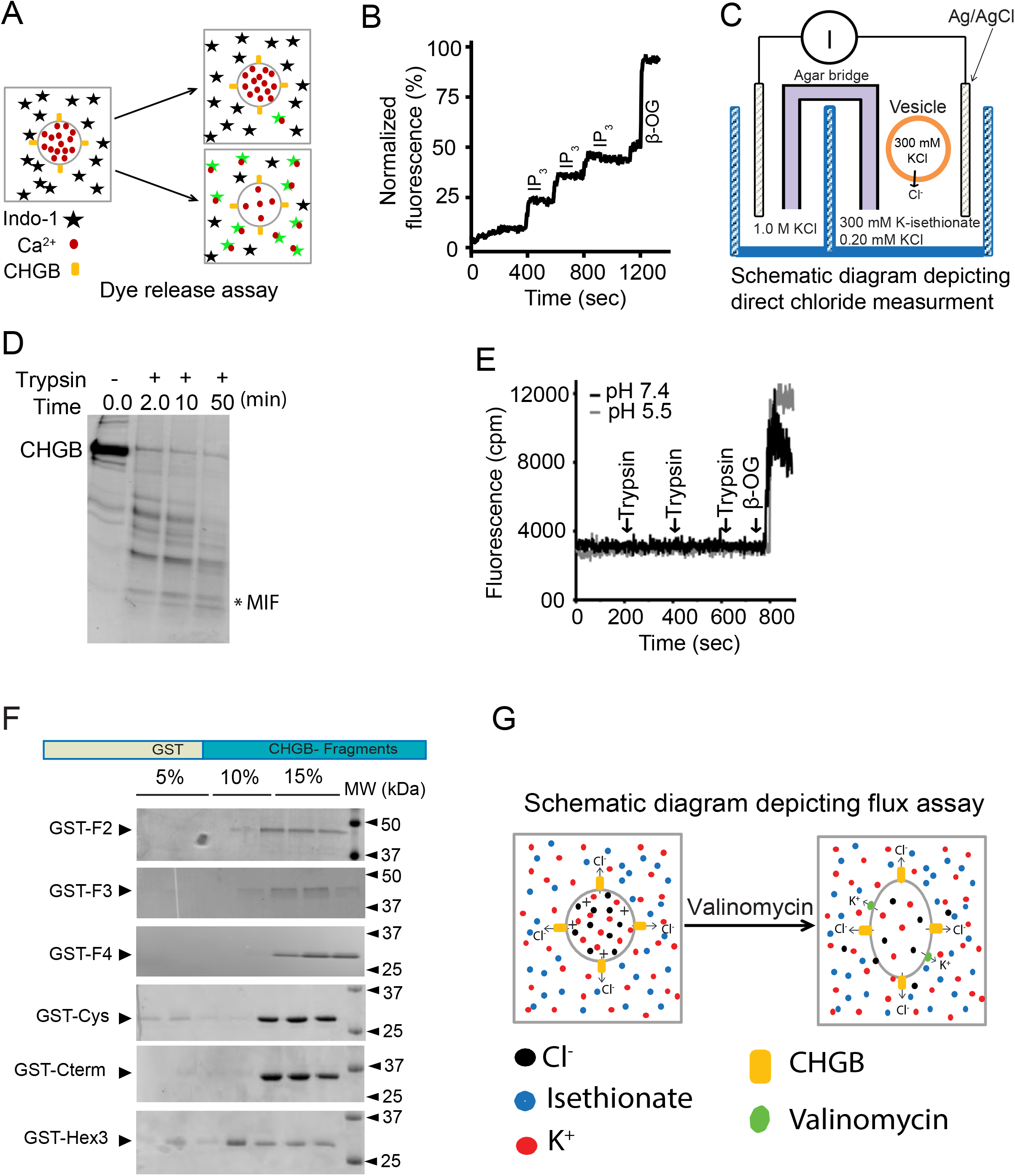
CHGB preferentially located to the outside of the vesicles and mapping its membrane interacting fragments. (**A**) Schematic diagram of the Ca^2+^-release assay. 1.0 micromolar Indo-1 (black star) in the outside would fluoresce stronger when Ca^2+^ ions (red dots) are released from vesicles and bound to Indo-1 (green stars). Excitation: 340 nm. Emission: 410 nm. (**B**) Positive control for the Ca^2+^ - release assay. Type 1 IP_3_R (a calcium release channel) protein was purified from rat cerebellum and reconstituted into egg PC vesicles with 1.0 mM CaCl_2_. The vesicles were changed into a Ca^2^+-free buffer before being used. The calcium release was induced by adding 1.0 micromolar IP_3_. Beta-OG was used to release all calcium at the end, and the data were normalized against the maximum signal. The experiments were repeated more than 4 times with similar results. (**C**). A diagram showing the direct measurement of chloride flux using a Ag/AgCl electrode. (**D**) 10 μg CHGB in egg PC vesicles treated with trypsin at RT. PMSF was used to stop trypsinization. Samples collected at various time points were separated in a 4-20% SDS-PAGE gel. The stable short fragment at ∼20 kDa was collected for mass spec analysis (marked as MIF), which is the CHGB-MIF. (**E**) Ca^2+^-loaded vesicles treated with trypsin while the fluorescence of Indo-1 was monitored as in **B**. The experiments were performed at pH 7.4 (black) and 5.5 (grey). (**F**) Coomassie-blue stained SDS-PAGE for the gradient fractions after the vesicle floatation experiments of different CHGB fragments as GST fusion proteins. (G) Schematic depiction of the light-scattering based flux assay.

**Supplementary Figure S4.**
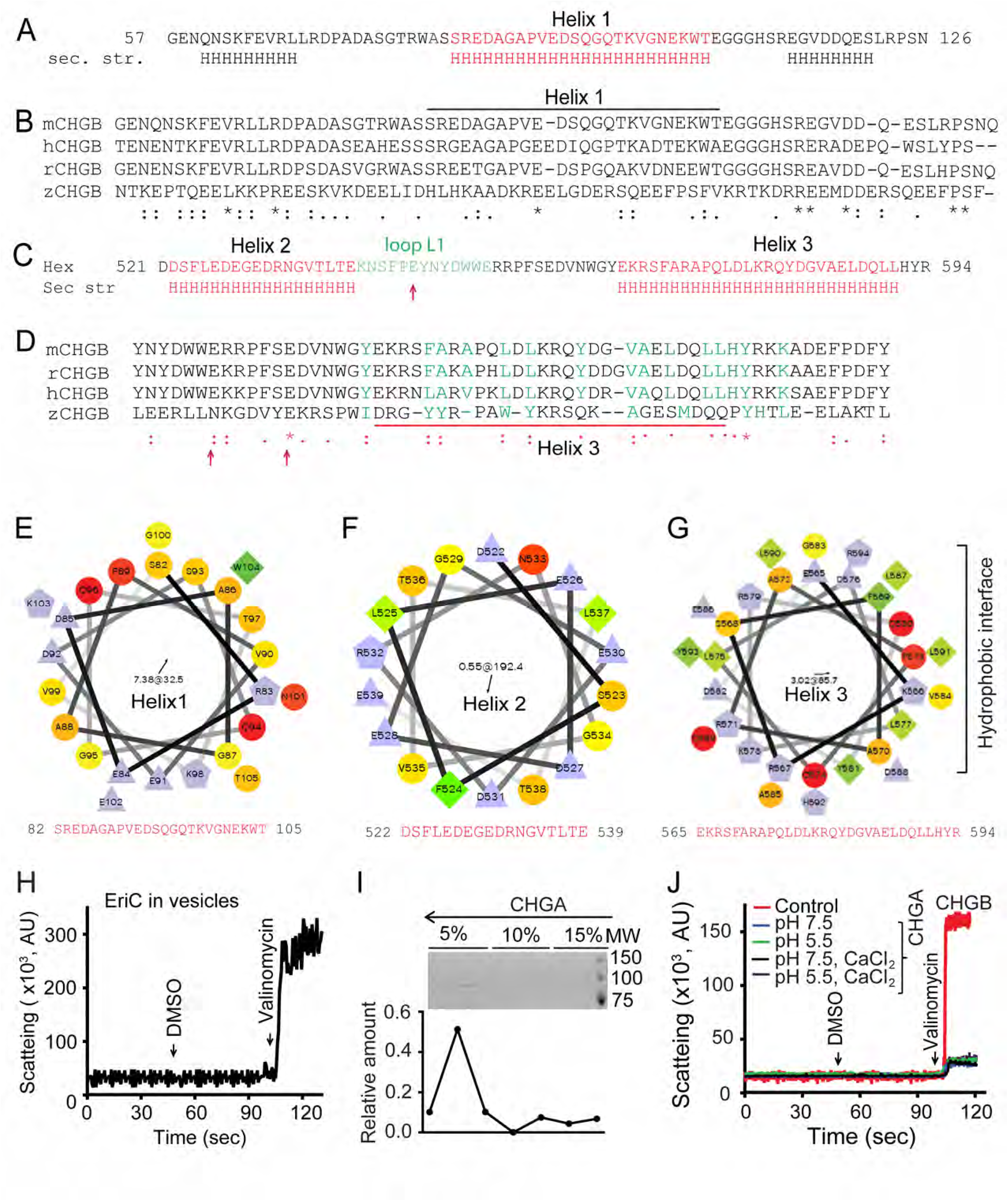
Structures of Helix 1 and the Hex fragment and control specimens for light-scattering-based flux assay. (**A**). Secondary structure prediction of the second half of the fragment F1 contains the Helix 1 (CHGB82-106). (**B**). Sequence alignment of the C-terminal half of the F1 fragments from different species (mCHGB: mouse CHGB; rCHGB: rat CHGB; hCHGB: human CHGB and zCHGB: zebrafish CHGB). (**C**). Secondary structure prediction of the Hex fragment (CHGB521-591) which contains two helical segments, Helix 2 and 3 (Hex 3) interspaced by a short loop. Hex 3 is the only long amphipathic helix in the MIF. (**D**) Alignment of the Hex 3 fragments from different species. The alignment was done using ClustalW and was manually adjusted. (**E-G**) Helical wheel representation of the CHGB helices 1-3 using an online server (http://rzlab.ucr.edu/scripts/wheel/wheel.cgi). (**H**) Light-scattering-based flux assay for EriC vesicles. EriC is a bacterial chloride/proton exchanger. (**I**) Vesicle floatation assay for the CHGA vesicles. Gradient fractions were run in SDS-PAGE and the protein distribution was quantified in the plot under the gel. (**J**) Light-scattering-based flux assay from CHGA vesicles in different extra-vesicular buffers (pH 7.5 and 5.5 with or without 1.0 mM CaCl_2_). The positive control of the CHGB vesicles (red trace) at pH 7.5 was showed for comparison.

**Supplementary Figure S5.**
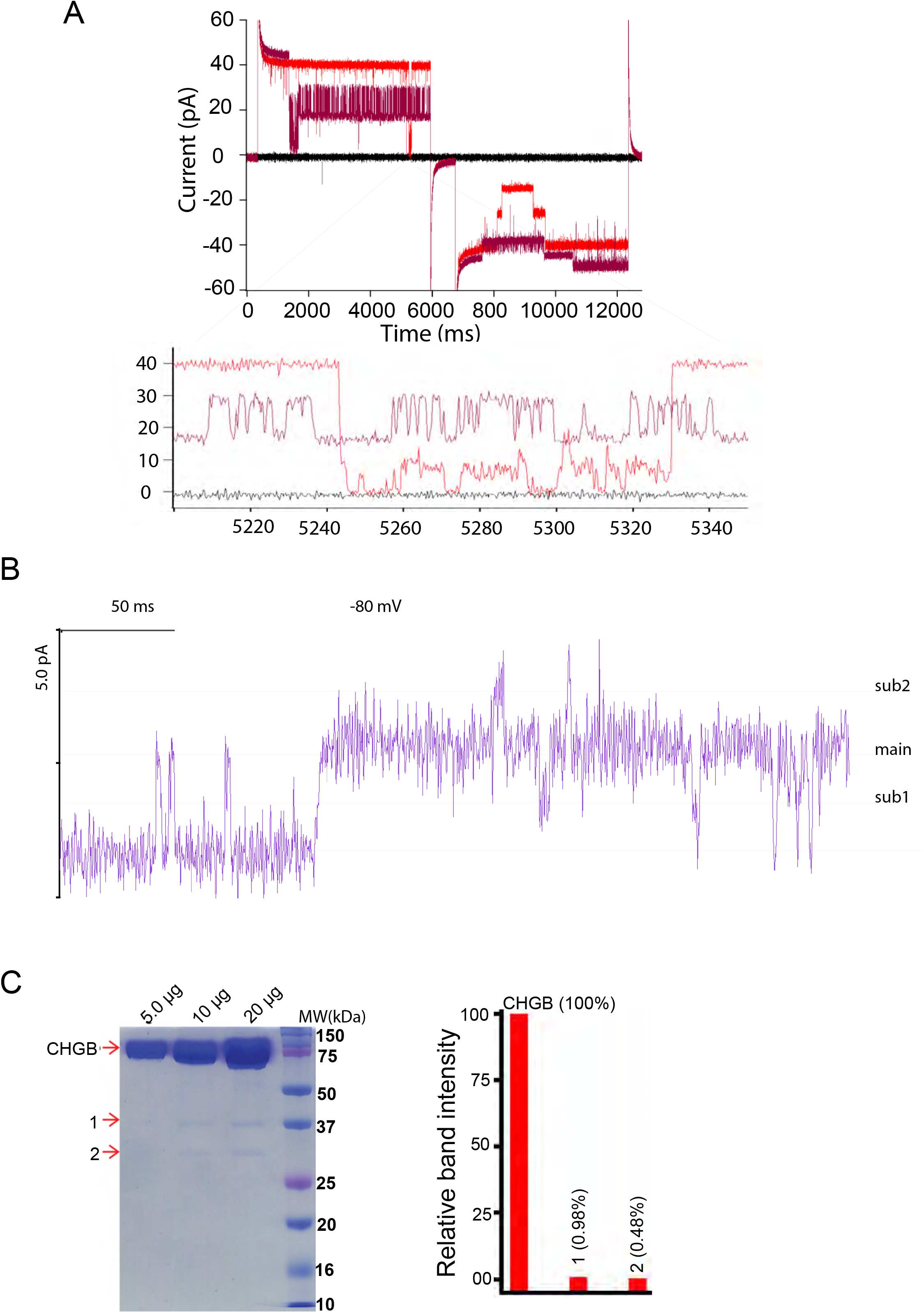
Bilayer recordings show little leak and reveal subconductance states. **A)**. The recordings from a multi-channel patch at ±140 mV, ±130 mV and 0 mV (dark) were showed to demonstrate that the recorded currents reached zero when all channels were closed, suggesting that the leaking currents were negligible and had very little effect in the recorded currents. The corresponding closing events were expanded in the bottom, which shows not only that the currents reached zero at multiple occasions, but also that there are subconductance states. **B)**. The recording at the −80 mV epoch was expanded to show the subconductance states (sub1 and sub2) besides the main state. **(C). Left**: 5, 10, 20 μg of purified recombinant CHGB were assayed in a Coomassie-blue-stained SDS-PAGE gel. The main CHGB band and the two contaminating bands (1 & 2) were cut out of the gels (at different times) and sent for mass spec analysis and proteomic identification. The main band was analyzed by the core-facility at UT Southwestern Medical Center and the two contaminating bands (1 & 2) were analyzed at UF ICBR. The identified protein candidates were listed in Supplementary Table 1 for the main band and Supplementary Table 2 for the two contaminating bands. **Right**: The scanned densities of these three bands were compared. The two contaminating bands account for only ∼1.4% of the density as the CHGB band, suggesting that the contaminants make a very small fraction of the total protein mass. From the data in table 3, the two contaminating bands are chiefly the partial degradation products of CHGB. These data together show that the amount of other proteins, if any, must be significantly less than 0.5% (the density of the contaminating band 2), and below 0.12%, which is the detection level in the gel. The purified CHGB is thus at least 99.8% pure with a small amount (< ∼1.5%) of degradation. In most cases, the degradation bands in freshly purified CHGB were not detectable as shown in Fig. 1B.

**Supplementary Figure S6.**
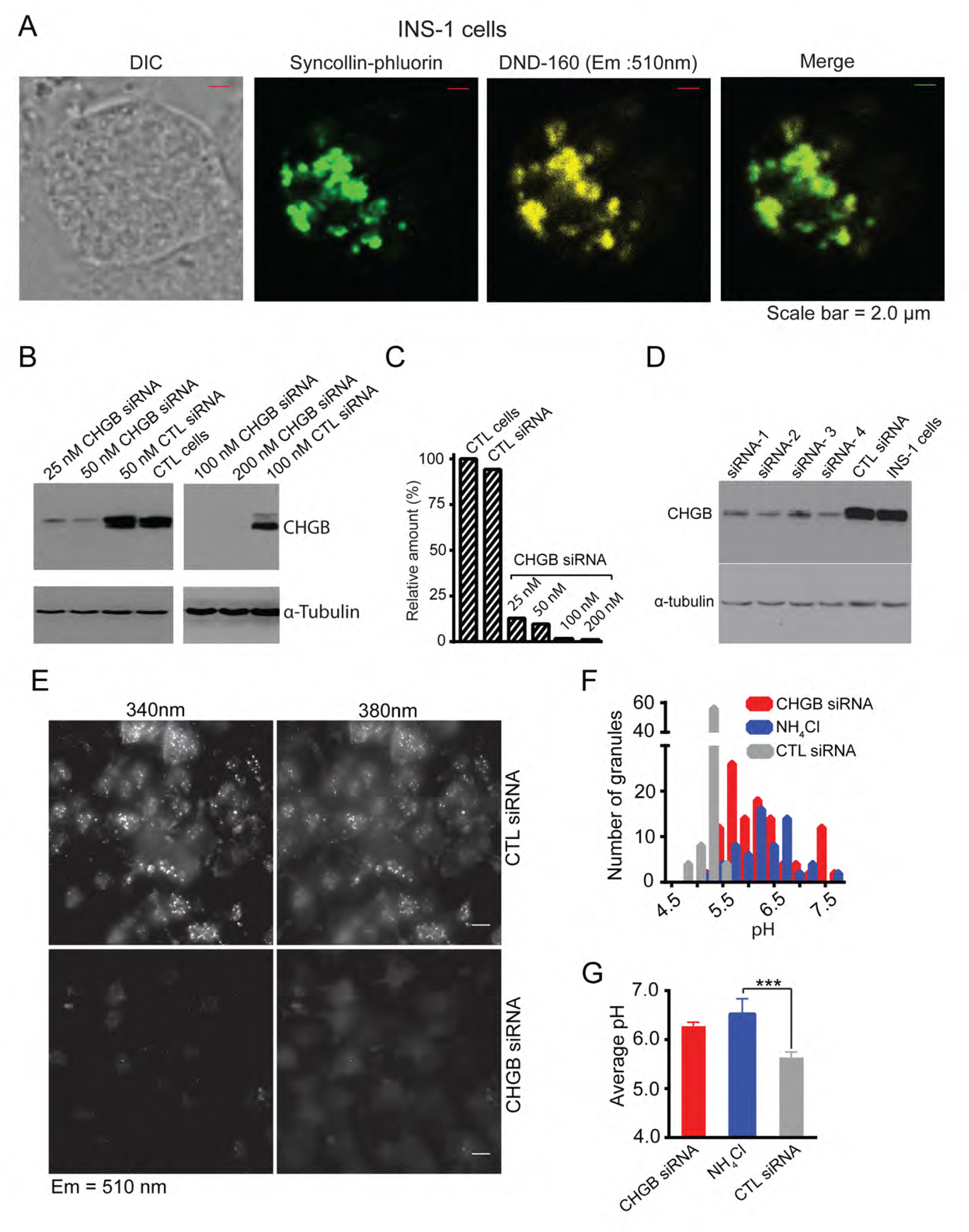
Establishment of intragranular pH measurement in INS-1 cells. **(A)** INS-1 cells expressing syncollin-pHluorin were stained with 0.5 μM DND-160 and imaged in a confocal microscope. DIC, syncollin-pHluorin and DND-160 images and the merge of the last two are shown to demonstrate the colocalization of syncollin and DND-160. **(B)** CHGB knockdown in INS-1 cell. Western blot of CHGB from cells transfected with different concentrations of CHGB siRNAs or control sequence-scrambled siRNAs (**scRNA**s), or control cells (CTL). ∼20 micrograms of protein from cell lysates were used for immunodetection. 100 nM siRNAs were used in other experiments. (**C**) Relative amount of CHGB protein from the same number of cells (based on the total protein and the loading control) under different conditions. (**D**) Four different siRNA molecules in the CHGB siRNA mixture were compared individually at 50 nM. Specific effects were seen for all four with the siRNA 2 and 4 more effective. The siRNA 2 and 4 were used separately or in combination to minimize off-target effects. (**E**) Dual excitation (340 and 380 nm) experiment for ratiometric measurement of intragranular pH. (**F**) Histogram showing the distribution of the granular pH read from the dual-excitation ratiometric measurements from images of INS-1 cells transfected with control siRNA (gray bars), CHGB siRNA (red bars) as well as cells treated with 5.0 mM ammonium chloride (blue bars). (**G**) Average pH values in the granules of three types of cells in **F**. The dual-excitation procedure followed what was reported before (Stienert P, et al., 2006).

**Supplementary Figure S7.**
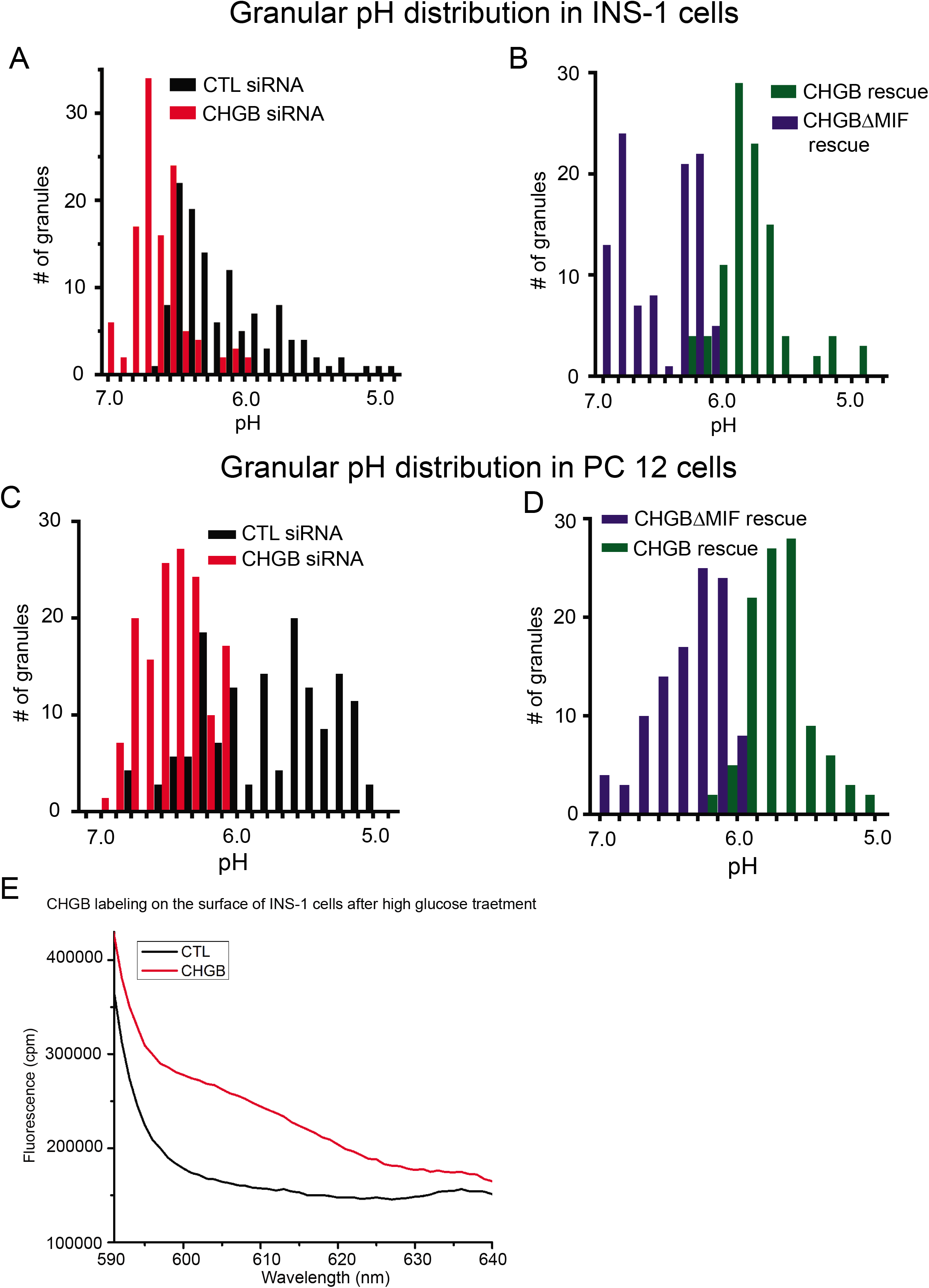
Distribution of ratiometric measurements in four differentially treated INS-1 cells and PC-12 cells. **(A)** Dual-emission measurements from INS-1 cells treated with control siRNAs (black) and CHGB siRNAs (red); and **(B)** Measurements from CHGB knockdown INS-1 cells transiently expressing CHGB (green, CHGB rescue) or CHGBΔMIF (blue). More than 100 granules were randomly picked from paired images of each type of cells. (C) Measurements from PC-12 cells treated with control siRNA(black) and CHGB siRNA (red); and **(D)** Measurements from CHGB knockdown PC-12 cells transiently expressing CHGB (green, CHGB rescue) or CHGBΔMIF (blue). More than 100 granules were analyzed for each type of cells. (E). CHGB was labeled with primary and secondary antibodies in the positive cells after glucose treatment. The control cells were not labeled with primary antibodies. The same number of cells in suspension was measured in a fluorometer to quantify the total labeling.

**Supplementary Figure S8.**
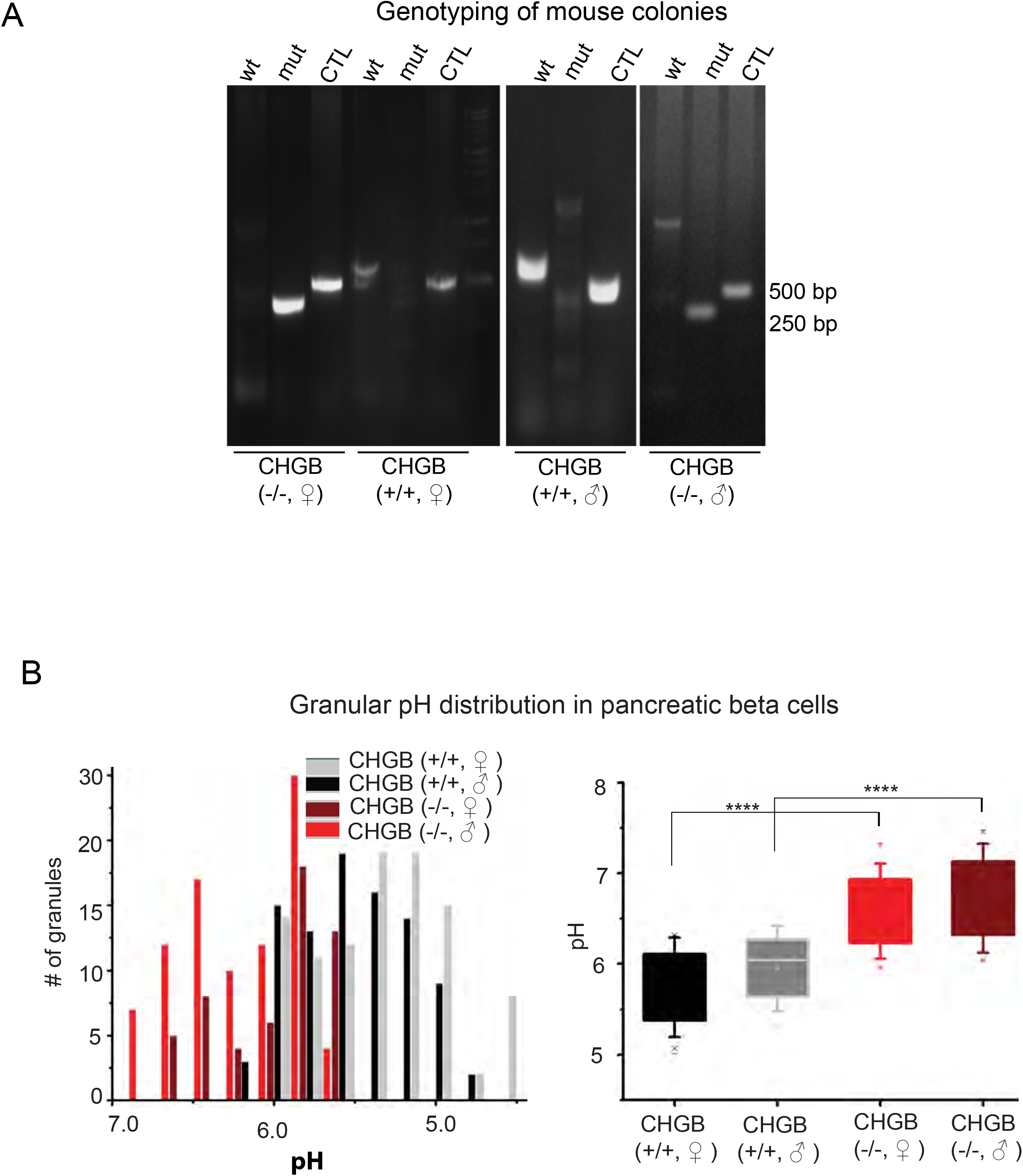
Characterization of the CHGB knockout mice. **(A)** Genotyping of mice. The wild-type and knockout alleles gave rise PCR fragments of different lengths. Homozygous male/female and wild-type male/female are compared here. (B). Left: Histograms of granular pH from beta-cells in four different types of mice. Control wild-type (male / female) mice are compared with homozygous CHGB knockout mice (male/female). Right: the average granular pH values are compared among the four different types of mice.

## Abbreviations

CHG: chromogranin
CHGB: chromogranin B
CHGB-MIF: a membrane interacting fragment at the C-terminal half of CHGB
ISG: immature secretory granules
DSG: dense-core granules
EM: electron microscopy
TGN: trans Golgi network
ClC: chloride channel
PC: phosphatidyl choline
RRG: readily releasable granule
SEC: size-exclusion chromatography
MIF: membrane-interacting fragment
SMCC: 4-(N-Maleimidomethyl) cyclohexane-1-carboxylic acid 3-sulfo-N-hydroxysuccinimide ester sodium

